# Divergent Outcomes Of Myeloid Cell Infection By Gammaherpesvirus 68 Reveal Stepwise Regulation By Host And Viral Factors

**DOI:** 10.1101/2023.06.21.545948

**Authors:** Gabrielle Vragel, Kyra S. Noell, Ashley Tseng, Rachael E. Kostelecky, Elizabeth A. Alexander, Shirli Cohen, Catalina Galvan, Manaal Dalwadi, Brittany D. Gomez, Eva M. Medina, Darby G. Oldenburg, Andrew Goodspeed, Eric T. Clambey, Linda F. van Dyk

## Abstract

The gammaherpesviruses establish a lifelong infection, with the cellular outcome of infection intimately regulated by cell type. Though myeloid cells are an early infection target, infection results in widely divergent outcomes without a clear explanation. Here we use murine gammaherpesvirus 68 to demonstrate that macrophages are readily infectable, resulting in three divergent infection outcomes dictated by viral and host factors. Infection in vivo and in a cell culture model results in a high frequency of viable cells characterized by restricted viral transcription and unexpected transcription of the ORF75 locus, with rare cells initiating but failing to complete lytic replication. Restricted infection can be fully converted to productive lytic replication by pre-treatment with the host cytokine IL-4. Finally, infection with a high viral dose also results in productive infection, but triggers increased cell death. These studies demonstrate stepwise regulation of virus infection and identify virus and host factors that dictate divergent outcomes in a single cell type.

## Introduction

The gammaherpesviruses (γHVs), including the human viruses Epstein-Barr Virus (EBV) and Kaposi’s sarcoma-associated herpesvirus (KSHV or HHV-8), are large double-stranded DNA tumor viruses. The γHVs establish a lifelong infection and are associated with chronic inflammation and cancer, particularly in immunosuppressed individuals [1–3]. Due to the strict host specificity of the human γHVs, murine gammaherpesvirus 68 (MHV68, γHV68 or MuHV-4) has been extensively used as a genetic and phenotypically relevant model to study γHV infection and pathogenesis in vivo [4, 5]. The γHVs infect multiple cell types, resulting in a range of outcomes including lytic infection characterized by active virus replication or quiescent latent infection [1], dictated by target cell type-specific virus-host interactions. Infection outcomes at the single-cell level can be heterogeneous, including abortive infection [6–10].

Myeloid cells, including macrophages, monocytes and dendritic cells are innate immune cells that have critical functions during γHV infection: they promote the early antiviral response yet are also targeted for infection and subject to viral manipulation. While there is clear evidence that KSHV, EBV and MHV68 infect myeloid cells [11–23], the state of infection in these cells remains ambiguous, with evidence for both lytic transcription [19, 24, 25] and latent infection [15, 18]; infection also appears to be abortive or unstable over time [26]. In some cases, myeloid cells are thought to facilitate transit to B cells where γHVs establish long-term latent infection [14, 26, 27]. γHV infection has also been shown to alter myeloid cell function, including KSHV-induced differentiation of dysfunctional macrophages [22], manipulation of innate immune signaling [16, 28, 29], and impaired function [30].

The most detailed insights regarding macrophage infection in vivo have come from the MHV68 system. MHV68 lytically replicates in macrophages prior to B cell infection [14, 27, 31]; MHV68 also establishes long-term latent infection in macrophages [15, 32]. While MHV68 passage through alveolar macrophages and marginal zone macrophages promotes establishment of B cell latency [14, 33], subcapsular sinus macrophages limit [34]. Macrophage infection is further regulated by viral (ORF4, ORF36 [24, 25]) and host factors (e.g. interleukin (IL)-4 and IL-13, Liver X receptors, cholesterol synthesis, type I and II interferons [35–38]).

Here, we sought to define γHV macrophage infection outcomes with a new level resolution using the MHV68 system. Early macrophage infection in vivo is dominated by cells with frequent transcription of the ORF75 locus, a γHV-conserved gene expressed in Kaposi’s sarcoma lesions and in EBV-transformed latently-infected B cells [39–42]. Using a reductionist system to provide robust, controlled and reproducible outcomes, we find that MHV68 robustly infects macrophages with three distinct outcomes regulated by virus and host factors: i) a restricted form of infection with rare cells undergoing lytic replication, ii) a transcriptionally active form of virus replication accompanied by high rates of cell death, and iii) fully productive lytic replication. These studies emphasize that macrophage infection is subject to multiple levels of control, resulting in diverse infection outcomes in a single cell type.

## RESULTS

### Single cell RNA sequencing analysis of acute MHV68 macrophage infection reveals two distinct transcriptional profiles

MHV68 macrophage infection has been associated with both lytic replication and latent infection [14, 15, 24, 32], potentially serving as an important intermediary prior to B-cell latency [14]. Despite these insights, the exact nature of myeloid cell infection remains incompletely characterized. To define the viral transcriptional signatures at the earliest stages of macrophage infection in vivo, we sort-purified MHV68 infected peritoneal macrophages from immunocompetent C57BL/6J mice at 16 hours post-infection, followed by single-cell RNA-sequencing (scRNA-seq). MHV68 infected cells were purified based on expression of a viral reporter gene encoded by the latency-associated nuclear antigen fused with the beta-lactamase enzyme (LANAβlac), a sensitive system that enables the direct identification and purification of both lytic- and latently-infected cells [43, 44]. As expected, Seurat-based clustering and UMAP-based visualization identified a dominant cluster of LANA+ peritoneal macrophages, with minor cell subsets including: i) macrophages with a proliferative gene signature (i.e. “macrophage proliferating”), ii) macrophages with an MHV68 lytic gene signature (i.e. “macrophage lytic”), defined by expression of two canonical lytic genes ORF7 and ORF54, which are abundantly expressed during lytic replication [45–47] and infrequent iii) B cells, iv) monocytes, v) dendritic cells and vi) T cells (Fig. 1A-C, Fig. S1A). Quality control metrics were relatively comparable across all cell subsets, including the “macrophage lytic” subset (Fig. S1B-E).

**Figure 1.**
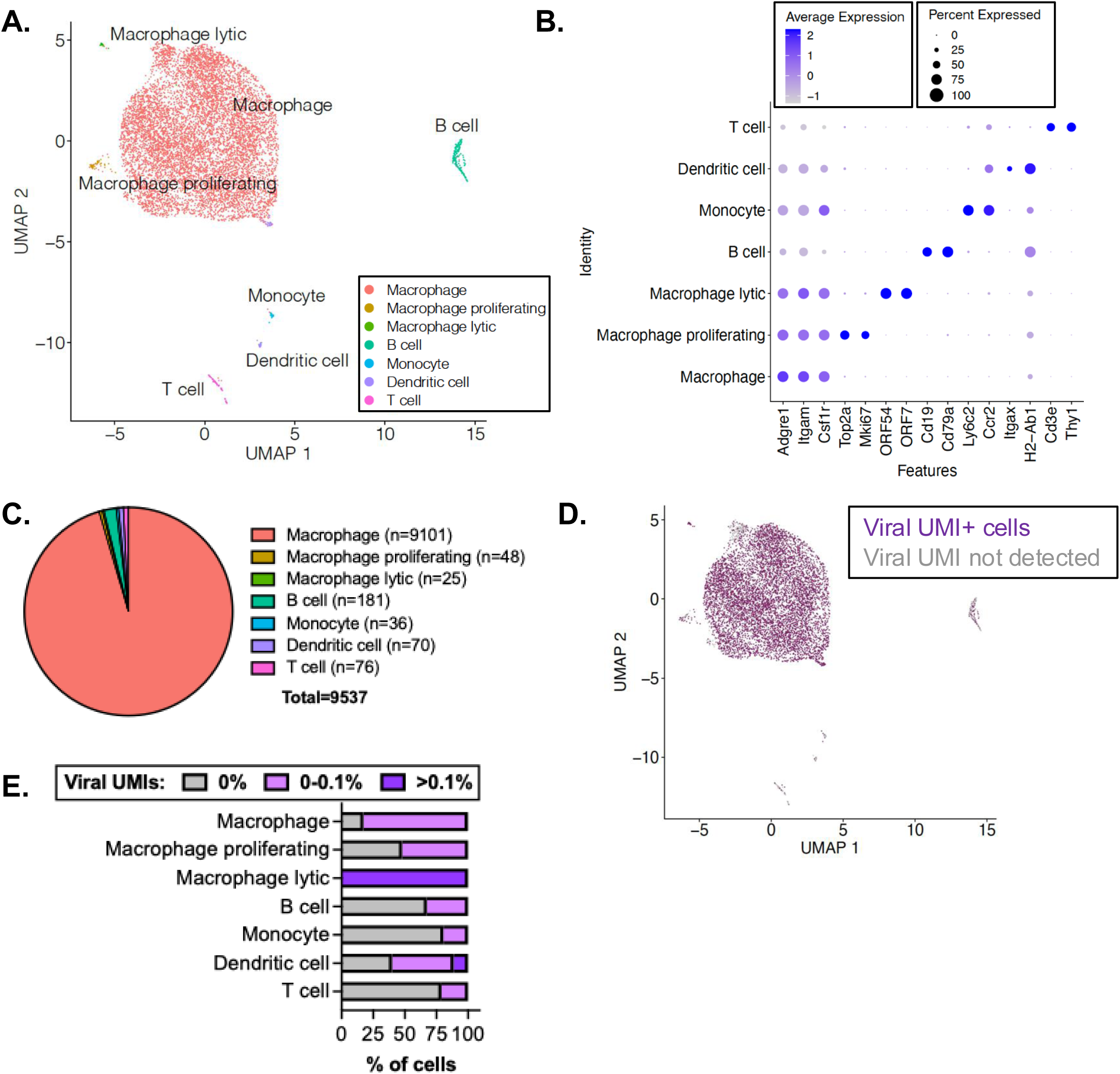
Single cell RNA sequencing analysis of acute MHV68 macrophage infection. C57BL/6J mice were infected with 1x10^6^ PFU of WT MHV68.LANAblac virus. MHV68-infected, LANAblac+ peritoneal cells were sort-purified at 16 hpi and subjected to scRNA-seq analysis. **(A-B)** Visualization of cell types based on lineage marker genes depicted in panel B. **(C)** Quantification of cell subsets defined by scRNA-seq. **(D)** UMAP visualization of cells with at least 1 viral UMI detected per cell identified by purple. **(E)** Frequency of cells stratified by the fraction of viral UMIs out of total UMIs detected per cell. 56 cells had >0.1% of total UMIs contributed by viral UMIs, including 23 of 9101 macrophages (0.25%), 25 of 25 macrophages with lytic gene expression (100%) and 8 of 70 dendritic cells (11.43%). Data are from 9,537 cells, using LANAβlac+ cells pooled from four infected mice analyzed as one sample in a single experiment, with cell subsets enumerated in panel C. Supporting data in **Fig S1-S2**.

When we visualized and quantified the frequency of cells with detectable viral unique molecular identifiers (UMIs), the majority (n=7,512; 82.54%) of LANA+ macrophages had detectable viral reads accounting for <0.1% of total UMIs per cell (Fig. 1D-E). In contrast rare macrophages with a lytic signature were characterized by a higher proportion (>0.1%) of viral reads per cell (Fig. 1D-E). The presence of cells with no detectable viral UMIs could either be due to the depth of sequencing or to the known sparseness of scRNA-seq data [48]. Increased granularity of cell clustering identified 14 cell clusters (Fig. S2) but did not significantly change defined cell populations nor the percentage of viral UMIs detected per cell subsets defined in Figure 1.

Next, we quantified the frequency of macrophages that expressed each viral gene, comparing the dominant macrophage population (n=9101 cells), with rare macrophages characterized by a proliferative (n=48 cells) or lytic (n=25 cells) gene expression signature. We detected ORF73-βlac mRNA, the transcript encoding the LANAβlac reporter by which cells were purified, in 12.89% of macrophages, with equivalent or higher frequencies of detection for only 3 other viral genes: ORF75C (30.75%), ORF75B (27.83%) and ORF75A (12.54%) (Fig. 2A). The apparent discrepancy between detection of LANAβlac protein and LANA transcript likely reflects the high sensitivity achieved by flow cytometry detection of enzymatic activity of the LANAβlac protein [43, 49], coupled with known limitations of scRNA-seq in detecting low abundance transcripts. Additional transcripts were detected in a lower frequency of cells: ORF72 (5.3%), ORF58 (3.5%), with detectable RNA for M3, ORF7, ORF50, ORF57, ORF61, ORF74, M12, M13, or M14 in 1-3% of cells (yellow circles, Fig. 2A). Remaining viral genes were only sporadically detected (<1% of cells). A similar hierarchy, albeit reduced in magnitude, was observed in proliferative macrophages (Fig. 2B). In contrast, rare lytic macrophages had a distinct viral gene signature: multiple lytic cycle-associated genes (ORF7, ORF54, ORF57, ORF58, and ORF61) were detected in 100% of cells (dark blue circles, Fig. 2C), with many more lytic cycle RNAs (M3, ORF6, ORF21, ORF38, ORF50, ORF52, ORF59, ORF72 and ORF75C) detected in >75% of cells (light blue circles, Fig. 2C). Detection of multiple, canonical lytic cycle RNAs in these cells is consistent with these cells initiating lytic gene transcription in vivo.

**Figure 2.**
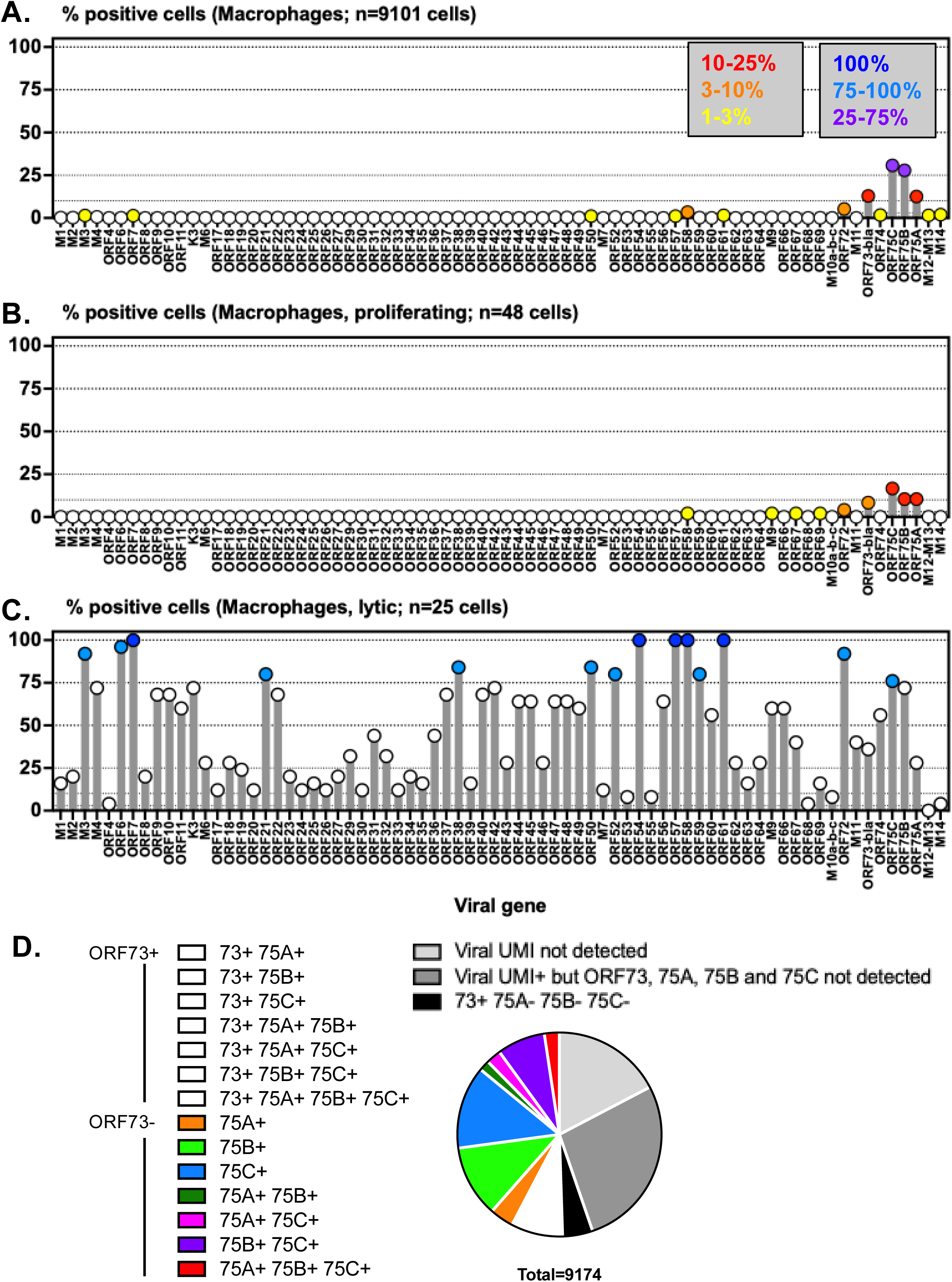
Single cell RNA sequencing analysis of acute MHV68 macrophage infection reveals two distinct transcriptional profiles. Analysis of viral gene expression defined by scRNA-seq in macrophage subsets from Figure 1**. (A-C)** Frequency of cells with detectable reads mapping to MHV68 genes, comparing (A) macrophages (82.79% viral UMI+), (B) macrophages with proliferative gene signature (52.08% viral UMI+), or (C) macrophages with lytic gene expression (100% viral UMI+). Viral genes are indicated on the x axis, arranged from left to right based on the published MHV68 genome, with the frequency of cells positive for each viral gene depicted on the y axis. To facilitate comparison of gene expression across the viral genome and cell populations, circles are color-coded to identify values that are in the indicated ranges (defined in key, panel A). Dotted lines indicate different thresholds of viral gene positivity, empirically defined. For panel C, only genes with >75% positivity were color-coded, given the limited number of cells and high viral positivity. **(D)** Frequency among macrophages defined by detection of ORF73, ORF75A, ORF75B, and ORF75C RNAs, enumerating all potential gene combinations for macrophages from panels A-C.

The ORF75 locus is conserved among γHVs, with MHV68 containing triplicated ORF75 genes, ORF75A, ORF75B and ORF75C [4, 50]. Previous studies of this locus have revealed multiple transcriptional units, including a polycistronic RNA that spans from the 5’ end of ORF75A through the 3’ end of ORF75C [50]. Though these genes likely arose from gene duplication, there is no significant similarity between the 3’ ends of ORF75A and ORF75B or ORF75A and ORF75C; only 22 of 26 nucleotides align between the final 150 nucleotides of ORF75B and ORF75C (defined by blastn [51], data not shown). The nucleotide divergence between ORF75 genes strongly suggests that UMIs mapping to each ORF75 gene were not due to mapping errors. Notably, 50.89% of LANAβlac+ macrophages had detectable expression of ORF75A, ORF75B and/or ORF75C (Fig. 2D). These studies suggest two distinct transcriptional signatures early after infection in vivo: 1) a high frequency of macrophages expressing transcripts from the ORF75 locus, 2) with rare cells characterized by lytic transcription.

### MHV68 readily infects J774 macrophages with limited lytic transcription

We next examined whether these viral outcomes could be observed in a reductionist cell culture model, assessing MHV68 infection in the murine myeloid-like cell line, J774, compared to 3T12 fibroblasts, a permissive murine fibroblast cell line [52]. As expected, MHV68 underwent robust lytic replication in 3T12s with extensive cytopathic effect (CPE) by 72 hours post infection (hpi). In contrast, MHV68-infected cultures of J774 cells showed minimal evidence of active virus replication, with no discernable CPE (Fig. 3A) and minimal viral genome replication or virus particle production above input virus (Fig. 3B-C). Infection with WT MHV68.LANAβlac resulted in a comparable frequency of LANAβlac+ cells in 3T12s fibroblasts and J774 cells (Fig. 3D-E, Fig. S3A), demonstrating that J774 cells are fully susceptible to MHV68 infection, initiating transcription and translation of ORF73 (an immediate early gene that is expressed in both latent and lytic infection, here detected using the LANAβlac fusion protein [43, 45–47]), with restricted infectious virus particle production.

**Figure 3.**
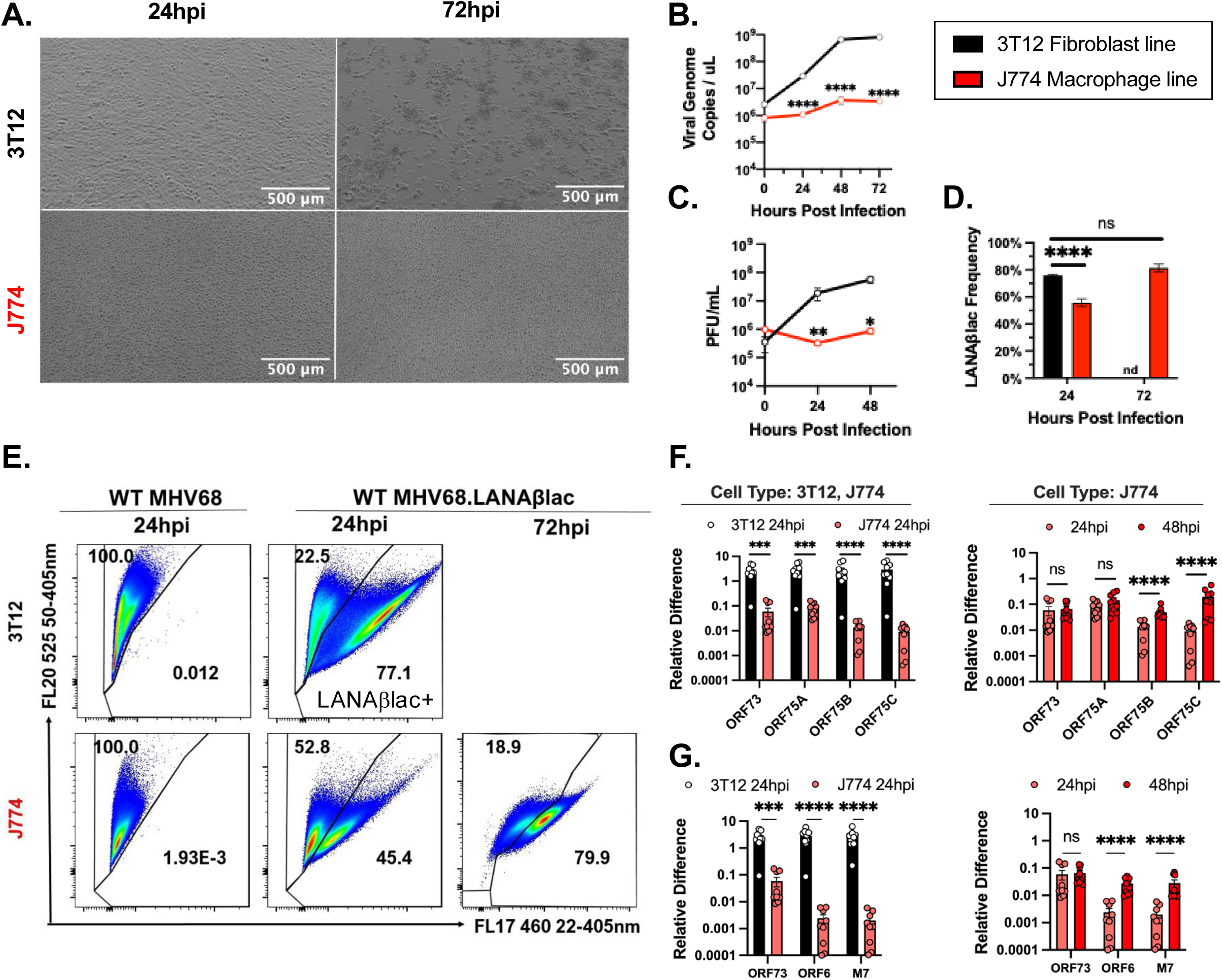
MHV68 readily infects J774 macrophages with limited lytic transcription. Analysis of MHV68 infection (WT MHV68.LANAblac, MOI=1 PFU/cell) comparing 3T12 fibroblasts and J774 macrophages. **(A)** Representative brightfield images of infected 3T12 (top) or J774 cells (bottom) early (24 hpi, left) or late (72 hpi, right) post-infection. 3T12 cells showed cytopathic effect (CPE), including cell rounding (24 hpi) and monolayer destruction (72 hpi), in contrast to J774 cells. **(B)** Viral DNA quantitation defined by qPCR. **(C)** Virus replication defined by plaque assay. **(D-E)** Frequency of MHV68-infected, LANAβlac+ cells defined by flow cytometry, detected by CCF2 substrate cleavage, analyzing single, viable cells. Samples infected with WT MHV68 (left, panel E) define background signal and do not encode LANAβlac. **(F-G)** Quantitation of viral gene expression defined by qRT-PCR, comparing 3T12 and J774 cells at 24 hpi (left) or J774 cells at 24 and 48 hpi (right). Flow cytometric analysis focused on single, viable cells. Data depict mean ± SEM, from 3 independent experiments with 3 biological replicates per experiment (B, D, F-G) or 2 (t=0) or 6 (t=24, 48 hpi) independent experiments (C), with individual symbols representing independent biological samples. Statistical analysis was done using unpaired t test, with statistically significant differences indicated, *p<0.05, **p<0.01, ***p<0.001, ****p<0.0001.

We next analyzed viral transcription in MHV68-infected J774 cells. ORF75A, ORF75B and ORF75C were readily detected in MHV68-infected J774 cells, albeit at lower levels than infected 3T12 cells (Fig. 3F, Fig. S3B). ORF75A expression was relatively comparable to ORF73, with minimal change in expression between 24 and 48 hpi, suggesting constitutive transcription of ORF73 and ORF75A (Fig. 3F). In contrast, ORF75B and ORF75C transcript levels were moderately lower than ORF73 at 16 and 24 hpi, with ∼4- and ∼22 fold increase between 24 and 48 hpi, respectively (right panel, Fig. 3F), potentially reflecting a subset of J774 cells initiating lytic infection. To assess lytic transcription, we quantified immediate early (IE) (ORF73), early/late (ORF6), and late gene (M7) expression [47]. ORF73 (and Blac) expression was reduced ∼10 fold in J774 myeloid cells relative to 3T12 fibroblasts (Fig. 3G, Fig. S3C-D), with ORF6 and M7 transcripts decreased >100 fold (Fig. 3G), consistent with greater deficits in lytic gene expression at later stages of the lytic cycle in MHV68-infected J774 macrophages.

ORF6 and M7 expression increased ∼10x between 24 hpi and 48 hpi (Fig. 3G, right panel), consistent with some lytic transcription in MHV68-infected J774 cultures.

### Only rare MHV68 infected macrophages display lytic gene expression

We next defined the frequency of cells with lytic gene expression using a series of flow cytometry-based assays. First, we infected 3T12 and J774 cells with a recombinant WT MHV68 virus, MHV68.HygroGFP (MHV68.H.GFP), that encodes a hygromycin resistance protein-GFP fusion under the transcriptional control of the human cytomegalovirus IE promoter inserted between ORF27 and 29b [53]. This virus robustly expresses GFP during active, lytic infection, identifying cells that initiated lytic transcription and translation. 3T12 cells had a much higher frequency of HygroGFP+ cells than J774 cells at 18 hpi (∼8.2% vs. ∼0.5% GFP+ respectively, Fig. 4A-B, Fig. S3A); a similar low frequency of HygroGFP+ cells was observed in both J774 and RAW264.7 macrophage cell lines (Fig. S4A). Infection with two additional reporter viruses indicative of lytic gene expression, encoding either an ORF59GFP gene fusion [54], or a viral cyclin-beta lactamase gene fusion (vCycβlac), further demonstrated that J774 cells had a greatly reduced frequency of lytic reporter gene positive events compared to 3T12 fibroblasts (Fig. 4C-D). The reduced frequency of lytic gene expression in J774 cells was further observed when we analyzed two proteins associated with lytic replication, the viral regulator of complement activation (vRCA, a direct measure of virus infection), a protein encoded by the ORF4 late gene, and phosphorylated histone H2AX (γH2AX, an indirect measure of infection), a host protein modification that occurs after DNA damage and during lytic replication as a direct target of the ORF36 viral protein kinase [24]. Whereas ∼45% of MHV68-infected 3T12 cells were vRCA+, γH2AX+, or vRCA+γH2AX+ by 24 hpi, the frequency of MHV68-infected J774 cells that expressed either of these proteins was significantly lower (∼2% cells at 24 hpi, ∼8% at 48 hpi) (Fig. 4E-F, Fig. S4B). Given the low frequency of cells expressing lytic cycle genes, we tested whether virally-infected J774 cells were capable of long-term growth and survival by purifying LANAβlac+ J774 cells. Over 14 days, LANAβlac+ J774 cells showed a progressive growth impairment relative to mock-infected cells, with limited evidence of viral replication, suggesting that the limited replication observed after MOI=1 infection in J774 cells is not simply due to delayed replication (Fig. S4C-I). These data indicate that MHV68-infected J774 cells rarely undergo lytic gene expression, mirroring in vivo findings, and suggest the utility of this system to interrogate MHV68-myeloid cell interactions.

**Figure 4.**
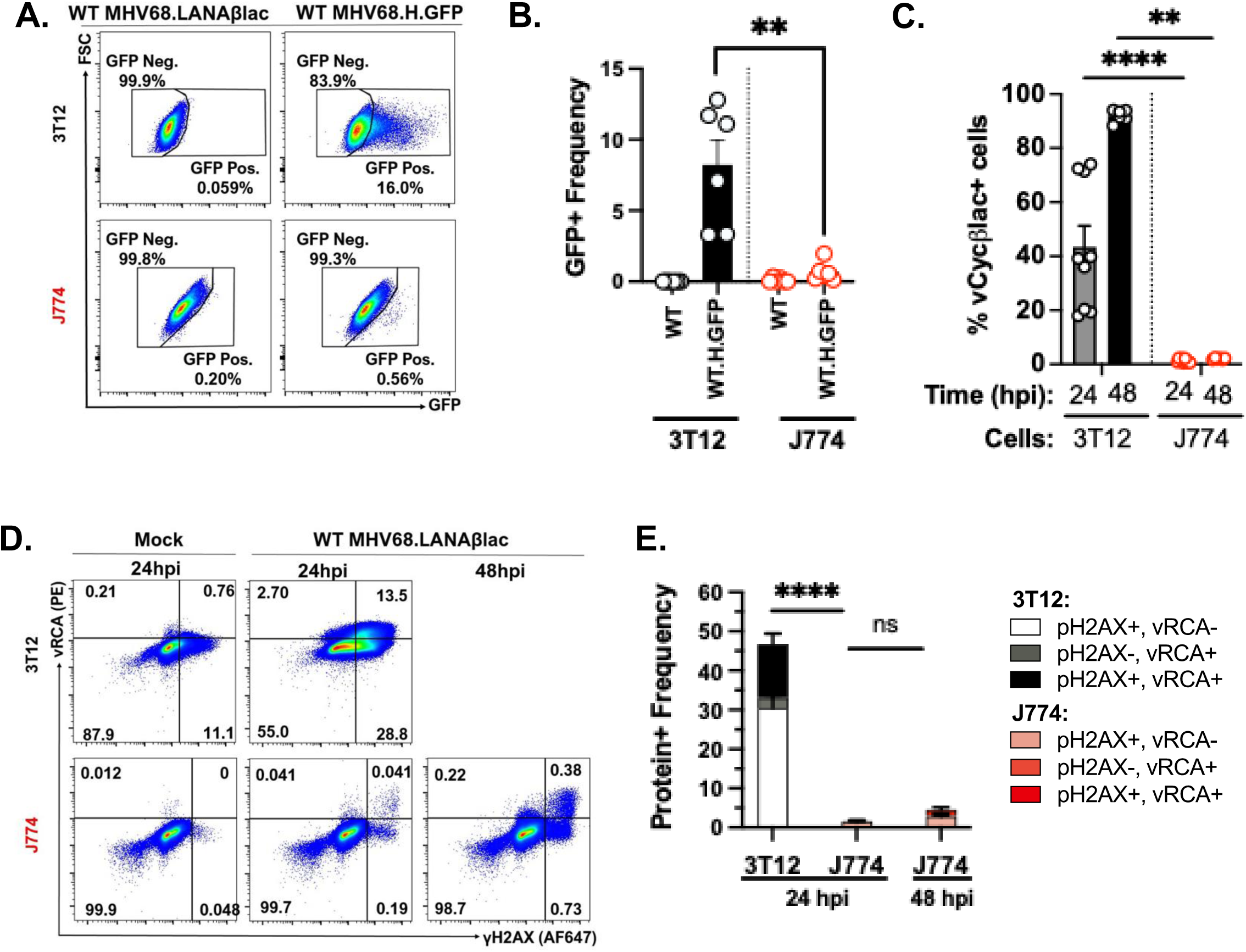
Only rare MHV68 infected macrophages display lytic gene expression. Analysis of MHV68 infection and lytic cycle profiles with either MHV68.H.GFP or MHV68.LANAblac (MOI=1 PFU/cell) comparing 3T12 fibroblasts and J774 macrophages. **(A)** Flow cytometric analysis of virus-expressed GFP in 3T12 or J774 cells 18 hpi. Data depict the frequency of cells with GFP expression, an indicator of MHV68.H.GFP infection. Background fluorescence defined by WT MHV68.LANAblac-infected cells (left column), without the addition of the CCF2 fluorescent substrate. **(B)** Frequency of HygroGFP+ cells defined by flow cytometry, as in panel A. **(C)** Frequency of vCycblac+ cells defined by flow cytometry. **(D)** Flow cytometric analysis of lytic infection-associated proteins, vRCA and γH2AX in 3T12 (top) or J774 (bottom) cells, comparing mock (left) or WT MHV68.LANAblac-infected cells. Data for 3T12 cells at 48 hpi was not assessed due to extensive cell destruction. **(E)** Quantification of the frequency of infected cells with protein expression associated with lytic infection defined by flow cytometry, as in panel D. Flow cytometry data focus on single, viable cells defined using gating strategies in **Fig. S3A, S4B**. Data show mean ± SEM from 3 experiments, with 2 (B,E) or 3 samples (C) per experiment. Statistical analysis done using Mann-Whitney test, with statistically significant differences as indicated, **p<0.01, ****p<0.0001.

### High dose MHV68 infection results in productive replication accompanied by a high frequency of cell death

Our studies to this point focused on infection using an intermediate infectious dose, a multiplicity of infection (MOI) of 1 plaque forming unit per cell. Since high MOI has been implicated in overcoming intrinsic antiviral immunity [55, 56], we next compared infection outcomes at an intermediate (MOI=1) or high inoculum (MOI=10). J774 cells infected with intermediate MOI had a low frequency of HygroGFP+ cells (∼4%); in contrast, infection with high MOI was characterized by a dramatically increased frequency of HygroGFP+ cells (∼92% cells) (Fig. 5A-B). Similar increases in lytic gene expression were observed following high-dose infection, quantifying: i) the frequency of ORF59GFP+ cells (Fig. 5C) and ii) the frequency of cells characterized by actin RNA degradation (Fig. 5D, an indirect measure of virus infection), a hallmark of virus-induced host shutoff mediated by the virally-encoded SOX exonuclease that occurs during lytic replication [6, 57]. These increased measures of lytic infection in high-dose cultures were characterized by increases in viral RNAs using bulk cultures (Fig. 5E), with high rates of cell death (Fig. 5F), evidence of viral protein expression (Fig. 5G-H) and production of new infectious virus (Fig. 5I). These data suggest that high-dose infection of J774 cells results in productive lytic infection, characterized by increased lytic gene expression, DNA replication and infectious virus production, in the presence of a high rate of cell death.

**Figure 5.**
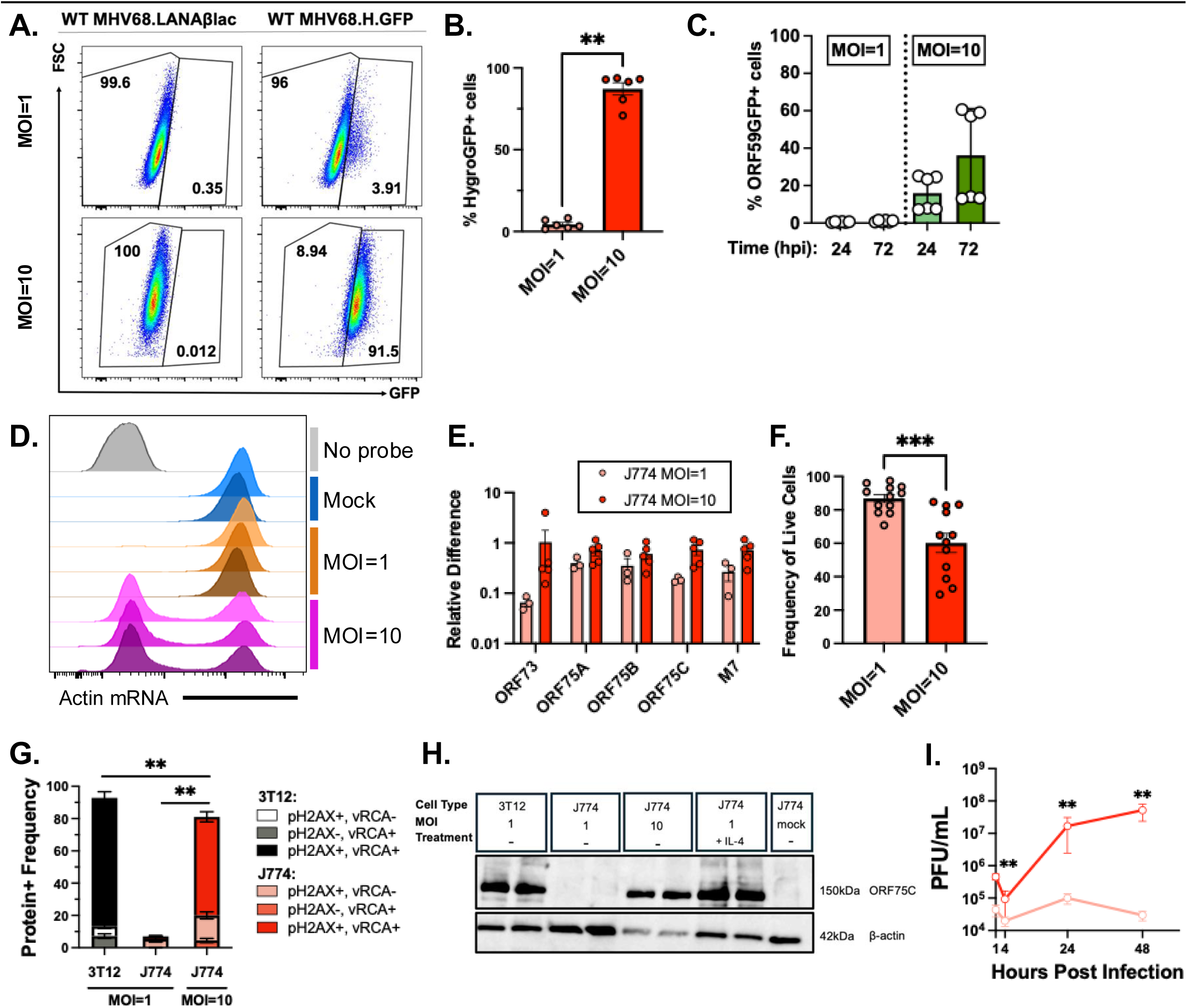
High dose MHV68 infection results in productive replication accompanied by a high frequency of cell death. MHV68 outcomes in J774 macrophages comparing intermediate (MOI=1 PFU/cell) or high dose (MOI=10) infection. **(A)** Flow cytometric analysis of GFP in MHV68.LANAblac- (left; negative control with no CCF2 substrate) or MHV68.H.GFP-infected cells (right) at MOI=1 (top) or MOI=10 (bottom). **(B)** Frequency of GFP+ cells quantified by flow cytometry, as in A. **(C)** Frequency of ORF59GFP+ J774 cells quantified by flow cytometry at 24 or 72 hpi. **(D)** Flow cytometric analysis of actin mRNA levels 24 hpi in mock or WT MHV68-infected cells with individual rows showing independent biological samples. **(E)** Quantitation of viral RNA expression by qRT-PCR standardized to 18S RNA, comparing intermediate or high MOI (MHV68.LANAblac) at the indicated timepoints. **(F)** Frequency of MHV68.HygroGFP-infected single cells that are viable, quantified by flow cytometry. **(G)** Quantification of the frequency of infected cells with protein expression associated with lytic infection defined by flow cytometry. **(H)** Western blot of viral ORF75C protein. Indicated cells were infected with MHV68.H.GFP or mock with or without 10ng/mL IL-4. Antibodies and molecular masses are indicated on the right. Blot shown is representative of 3 independent experiments**. (I)** Virus replication defined by plaque assay. Flow cytometry data analyzed single, viable cells (A-D, G) or single cells (F), with gating strategy in **Fig. S3**. Data show mean ± SEM, with individual symbols showing independent biological samples from 2-3 independent experiments with 2-3 biological replicates per experiment. Statistical analysis was done using Mann-Whitney test with statistically significant differences indicated, **p<0.01, ***p<0.001.

### MHV68 infection of macrophages is modulated by viral and host factors

Given that interleukin (IL)-4 induces viral replication and reactivation from latency in MHV68-infected primary macrophages [37, 58], we tested whether IL-4 pre-treatment was sufficient to overcome the blocks on lytic replication following infection with an intermediate dose. J774 cells that were pre-treated with IL-4 prior to infection at MOI=1 had a greatly enhanced frequency of HygroGFP+ cells with ∼60% GFP+ cells by 48 hpi (Fig. 6A-B), resulting in a time- and concentration-dependent increase in viral DNA replication relative to untreated cells (Fig. 6C), culminating in increased infectious virus production (Fig. 6D). These data demonstrate that J774 cells are fully competent to support MHV68 lytic replication, evidenced by IL-4 pre-treatment overcoming restricted MHV68 replication in J774 macrophages, consistent with the effects of IL-4 on in vivo macrophage reactivation [55]. To begin to define how viral factors regulate macrophage infection, we examined two viral genes: the viral cyclin (vCyc), dispensable for lytic replication in fibroblasts but critical for reactivation from latency [59], and the immediate early transactivator Rta, which is essential for lytic replication and capable of initiating reactivation from latency [60–62]. vCyc-deficient MHV68 infection was associated with a greatly reduced frequency of cells that expressed either vRCA and/or γH2AX relative to WT MHV68-infected cells (Fig. 6E). Conversely, infection with a virus that overexpresses Rta (CA-RTA MHV68) was associated with a greatly increased frequency of cells expressing these lytic cycle indicators (Fig. 6E). These data demonstrate that the vCyc is necessary to promote lytic gene expression in macrophages and that Rta is sufficient to drive lytic gene expression in macrophages, illustrating how host and viral factors shape the outcome of macrophage infection.

**Figure 6.**
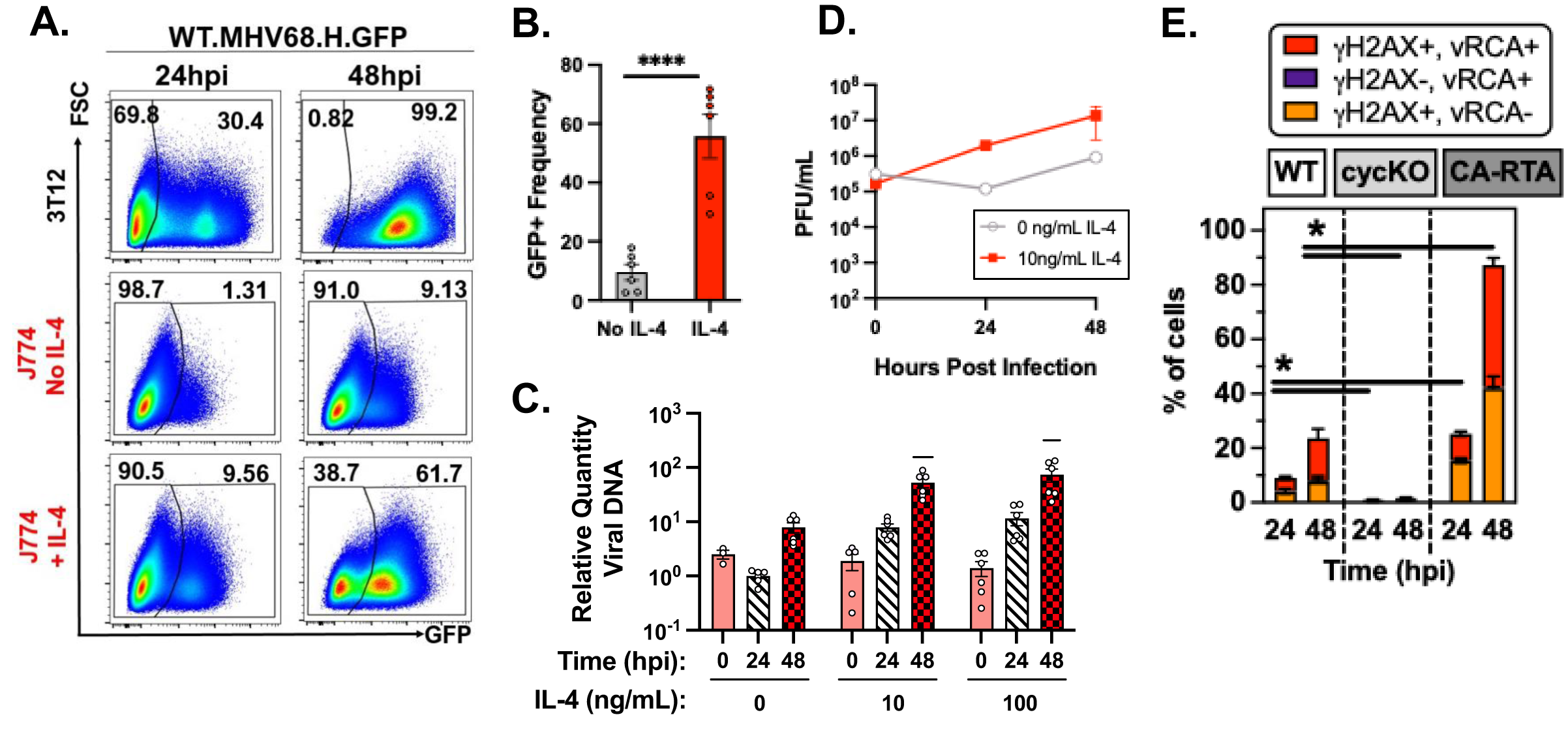
MHV68 infection of macrophages is modulated by host and viral factors. All infections done with MOI=1. **(A-D)** IL-4 effect on MHV68 infection and genome replication in J774 macrophages. IL-4 cultures were pretreated with IL-4 for 16-18 hours at the indicated concentration, with fresh IL-4 replenished after infection. **(A)** Flow cytometric analysis of MHV68.H.GFP infection in 3T12 (top), untreated J774 (middle) or IL4-treated J774 cells at 24 (left) or 48 hpi (right), analyzing single, viable cells. **(B)** Frequency of GFP+ cells quantified by flow cytometry, as in A. **(C-D)** IL-4 effect on viral DNA replication **(C)** and infectious virus production **(D)** in WT MHV68-infected J774 cells. **(E)** Frequency of J774 cells expressing markers of lytic infection (γH2AX, vRCA) quantified by flow cytometry, comparing WT, vCycKO or CA-RTA MHV68 at 24 or 48 hpi. Flow cytometric analysis focused on single, viable cells. Data depict mean ± SEM, with individual symbols representing independent biological samples from 2-3 independent experiments with 2-3 biological replicates per experiment (B,C,E) or from 1-2 independent experiments with 3 biological replicates measured (D). Statistical analysis used (B) unpaired t test, (C) one-way ANOVA comparing samples collected at the same timepoint untreated versus cytokine treated, or (E) one-way ANOVA comparing samples collected at the same timepoint with WT infection (nonparametric Kruskal-Willis test subjected to Dunn’s multiple comparison test), with statistically significant differences indicated, *p<0.05, **p<0.01, ****p<0.0001.

## DISCUSSION

Myeloid cells, including macrophages, monocytes and dendritic cells, are well-documented targets of γHV infection [11–27], though there remains little consensus on how infection is regulated in these cells or why infection results in widely divergent outcomes. Here, we present an in-depth analysis of MHV68 infection of macrophages, documenting multi-layered regulation of outcomes. First, macrophages harvested early after MHV68 infection in vivo are characterized by frequent transcription of the ORF75 locus in the absence of other detectable viral RNAs. Second, MHV68 induces three divergent outcomes in a reductionist system of macrophage infection: i) a restricted state of infection, reminiscent of latency, with infrequent cells initiating lytic infection, ii) a productive and destructive form of lytic replication associated with a high frequency cell death or iii) full lytic replication. Third, divergent infection outcomes are readily observed by manipulating virus (multiplicity of infection, viral mutations) and host factors (cytokine exposure), illustrating how divergent infection outcomes can occur in a single cell type. These data suggest that MHV68 replication in macrophages is blocked in early transcription that is overcome by IL-4 pre-treatment (Fig 4), and that high MOI infection results in productive infection that is marked by cell death. These studies further suggest a previously unappreciated role for the ORF75 locus as a potentially important regulator of macrophage infection, and suggest re-evaluation of previous assumptions (e.g. that macrophages poorly support lytic replication).

By examining the heterogeneity of macrophage infection at single cell resolution, our studies provide a refined perspective of MHV68 infection in macrophages. These studies unequivocally identify that MHV68 robustly infects macrophages and are capable of initiating transcription and translation of at least the ORF73 gene, with only a minor fraction of these cells showing hallmarks of lytic replication. Whether the majority of LANA+ cells reflect a bona fide latent population that meets the formal definition of herpesvirus latency or a previously undescribed, restricted form of infection remains an open question. The formal definition of herpesvirus latency includes virus genome retention, limited gene expression and the capacity to resume lytic gene expression at minimum, with evidence of a circularized genome in the absence of linear genomes often used as an additional criterion. We therefore refer to this macrophage infection as ‘restricted’ to accurately reflect that they are virus genome positive and limited in virus gene expression, yet it remains to be seen whether they fully meet a formal definition of latency. Although selective ORF75 transcription has not been reported in the context of MHV68 macrophage infection, transcripts mapping to the ORF75 locus have been observed in some in vitro latency models, a finding previously attributed to potential detection of unspliced ORF73 transcripts [45, 63, 64]. Here, we detected individual ORF75 gene products (i.e. ORF75A, ORF75B, and ORF75C) by polyA-based scRNA-seq, strongly suggesting that these RNAs represent ORF75-specific transcripts, not unspliced ORF73 RNA. These findings further suggest that ORF75 transcription is regulated in additional ways beyond those observed for lytic replication [50]. Notably, ORF75 transcription has been reported in contexts beyond lytic replication for EBV BNRF1 (the ORF75 ortholog), where it is expressed in lymphoblastoid cell lines [41], and for KSHV ORF75, where it is detected in Kaposi’s sarcoma lesions [65, 66].

Recent studies have shown that the KSHV ORF75 promoter is regulated in a cell type-specific manner, active in endothelial cells but repressed in B cells [40]. Collectively, these findings suggest that the ORF75 locus may be subject to macrophage-specific regulation, that ORF75 gene products may have unique functions in macrophages, and that ORF75-targeted interventions (e.g. vaccination against ORF75-derived T cell epitopes [67–69]) may selectively disrupt macrophage infection.

Through use of a reductionist system of macrophage infection, we find that MHV68 infection can result in three divergent outcomes. Using infection with an intermediate infectious dose (MOI=1), MHV68 readily infects the majority of cells, defined by LANAβlac expression, with only a minor fraction of cells characterized by lytic protein expression and no measurable viral DNA replication or infectious virus production; this finding parallels our in vivo analysis. While we can detect LANAβlac+ J774 cells for up to 14 dpi, whether this in vitro model represents bona fide latency remains unknown at this time. In striking contrast to intermediate dose infection, infection with a high virus dose (MOI=10) greatly increases the frequency of cells undergoing productive lytic infection, but is accompanied by a high rate of cell death. These outcomes suggest that macrophage infection is subject to multiple levels of regulation: i) a block that limits entry into the lytic cycle (observed with an intermediate infection dose) that can be overcome by a high MOI, with ii) profound activation of cell death pathways upon infection with a high MOI. The molecular mechanisms underlying this cell death, and what impact this process has on virus replication remains an active area of investigation. Despite these barriers to lytic replication, it is notable that pre-treatment with the cytokine IL-4 induces a high frequency of lytic transcription, viral DNA replication and infectious virus production, consistent with previous reports that helminth infection and IL-4 can promote virus replication in macrophages through STAT6-dependent transcriptional activation of RTA, the master lytic viral transactivator [37, 58].

By investigating the interplay between viral and host factors in macrophage infection, our studies demonstrate significant heterogeneity in infection outcome in macrophages. These findings provide an important new perspective on previously published findings and help to resolve previously ambiguities in the literature. First, the magnitude of MHV68 lytic replication within bone marrow macrophages or myeloid cells is relatively modest compared to fibroblasts [25, 70], with lytic antigen detected in a subset of bone marrow-derived macrophages and RAW264.7 macrophages [30, 37, 58]. Our data are consistent with this and emphasize that cells initiating lytic transcription represents only a fraction of infected cells. Second, there are numerous studies that have identified positive and negative regulators of macrophage infection. MHV68 lytic replication in macrophages is supported by multiple viral proteins (viral protein kinase [24], vRCA [25], vCyclin, Rta) and host factors (H2AX [38], ATM [71] and apolipoprotein E [72]). Parasitic infection can further promote macrophage infection through two distinct mechanisms, either by eliciting IL-4 or IL-13, STAT6-inducing cytokines that directly transactivate the MHV68 gene 50 promoter [37], or by expanding the number of large peritoneal macrophages allowing for an increased frequency of infection targets in vivo [73, 74]. The capacity of IL-4 to alter primary infection of J774 macrophages expands the potential impact of cytokine responses and co-infections and their potential to alter the outcome of primary infection. Conversely, MHV68 replication in macrophages is limited by type I and type II interferons [75], IRF1 [76], HDAC1 and 2 [77], liver X receptors [35], and Low-Density Lipoprotein Receptor [36] and chromatin accessibility [75], with complex regulation by innate sensing [28] and cell cycle and DNA damage signaling [38]. In vivo, IFNγ constrains MHV68 reactivation from latency within peritoneal macrophages [78] and CD4 T cells constrain chronic macrophage infection in the lung [70]. Based on our current findings, we postulate that these diverse regulatory mechanisms may affect the relative proportion of cells initiating lytic replication with MHV68, or may fundamentally alter the susceptibility of infection across all cells. Defining how these regulatory mechanisms influence the frequency and outcome of macrophage infection at the single-cell level remains an important future goal.

This study has important limitations. First, our studies of primary in vivo infection focus on a single, early timepoint that does not allow extrapolation to the impact on the dynamics of infection which follow. Conservatively, we can conclude that the high frequency of LANA+ macrophages with detectable ORF75 transcripts is not explained by the current paradigm of lytic or latent transcription (e.g. exemplified by [47]) and suggests additional mechanisms of transcriptional regulation within the ORF75 locus beyond that in fibroblasts [50]. Second, our in vitro studies primarily rely on the J774 mouse myeloid cell line, a reductionist in vitro system to study MHV68-macrophage interactions. Though these cells demonstrate parallels with primary macrophages (e.g. infrequent lytic replication at baseline that can be potently enhanced by IL-4 pretreatment), whether the precise molecular mechanisms engaged in these cells reflect one or more primary macrophage subsets remains to be determined. This is an important, but challenging question, given that macrophage subsets in vivo are known to respond differently to infection (e.g. large versus small peritoneal macrophages differ in their infection frequency with MHV68 early after infection [73, 74]; infection of marginal zone macrophages is thought to precede B cell latency [14, 33], whereas subcapsular sinus macrophages limit infection [34]). Despite these limitations, our results are consistent with previous studies demonstrating lytic protein expression in a subset of bone marrow-derived macrophages and RAW264.7 macrophages [30, 37, 58], and revealing that high MOI can overcome genetic requirements for lytic gene expression at a lower MOI [38]. Finally, it is important to recognize that the outcome of γHV-myeloid cell infection may be affected by the mode of virus entry. While our experiments test macrophage infection using cell-free virus, additional modes of infection have been reported (i.e. virus internalization via epithelial cell presentation [31], or infection through an Fc receptor-dependent mechanism [30]). These factors would likely be influenced by the route and stage of infection and host immune status.

In total, our studies provide critical new insights into how γHV infection of myeloid cells is regulated, demonstrating unanticipated transcription during primary infection in vivo and three divergent infection outcomes which can be readily manipulated by virus and host factors. Our studies highlight how bulk analyses and subsequent conclusions drawn from these analyses are insufficient to describe the heterogeneity of MHV68 infection in myeloid cells – a conclusion that could apply to γHV interactions beyond the myeloid compartment. These data redefine a conceptual framework to study myeloid cells as a γHV target and emphasize the knowable complexity of γHV-myeloid cell interactions.

## STAR METHODS

| REAGENT or RESOURCE | SOURCE | IDENTIFIER |
| --- | --- | --- |
| **Antibodies** |  |  |
| Anti-phospho-histone H2A.X (Ser139), AF647 conjugate | Sigma Aldrich | Cat# 07-164-AF647<br>RRID: AB_11213838 |
| Rabbit anti-ORF4 | Kapadia et al.<sup>[79]</sup> | NA |
| Zenon Rabbit IgG Labeling Kit, R-PE | Thermo Fisher Scientific | Cat# Z25355 |
| F4/80-AlexaFluor 674 Clone BM8 | Thermo Fisher Scientific | Cat# MF48021<br>RRID: AB_10375289 |
| Anti-mouse CD16/CD32 (Fc Shield) (2.4G2) | Tonbo Biosciences | Cat# 70-0161 |
| chicken anti-ORF75C | Santana et al.<sup>[80]</sup> | NA |
| mouse monoclonal B-actin | Thermo Fisher Scientific | Cat# MA5-15739,<br>RRID: AB_10979409 |
| **Bacterial and virus strains** |  |  |
| Virus: WT MHV68 | Virgin et al.<sup>[4]</sup> | NA |
| Virus: WT MHV68.LANA::blac | Nealy et al.<sup>[43]</sup> | NA |
| Virus: MHV68.HygroGFP | Forrest and Speck et al.<sup>[53]</sup> | NA |
| Virus: MHV68.ORF59.GFP | Smith et al.<sup>[54]</sup> | NA |
| Virus: MHV68.vCycKO | Van Dyk, Virgin, and Speck et al.<sup>[59]</sup> | NA |
| Virus: MHV68.vCyc::blac | Dr. Darby Oldenburg, unpublished data | NA |
| Virus: MHV68.CA-RTA | Rickabaugh et al.<sup>[81]</sup> | NA |
| **Chemicals, peptides, and recombinant proteins** |  |  |
| Mouse IL-4 | PeproTech | Cat#214-14 |
| Trypan Blue Dye | Bio-Rad | Cat# 1450021 |
| CCF2-AM | Thermo Fisher Scientific | Cat# K1023 |
| LIVE/DEAD Fixable Blue | Thermo Fisher Scientific | Cat# L23105 |
| LIVE/DEAD Fixable Near IR | Thermo Fisher Scientific | Cat# L10119 |
| **Critical commercial assays** |  |  |
| Chromium 3' V3 Library Kit | 10x Genomics | NA |
| OneStep RT-PCR kit | Qiagen | Cat# 210212 |
| LightCycler 480 Probe Master Mix Kit | Roche | Cat# 04707494001 |
| Quantitect Primer Assay | Qiagen | Cat# 249900 |
| PrimeFlow RNA Assay Kit | Thermo Fisher Scientific | Cat# 88-18005-210 |
| DNeasy Blood and Tissue kit | Qiagen | Cat# 69504 |
| **Deposited data** |  |  |
| Single Cell RNA-seq | This paper | GSE233202 |
| **Experimental models: Cell lines** |  |  |
| BALB/3T12-3 | ATCC | Cat# CCL-164 |
| J774A.1 | ATCC | Cat# TIB-67 |
| RAW 264.7 | ATCC | Cat# TIB-71 |
| **Experimental models: Organisms/strains** |  |  |
| Mouse: C57BL/6 mice (8 weeks) | The Jackson Laboratory | #000664 |
| **Oligonucleotides** |  |  |
| qPCR, RT-qPCR, PCR Primers | This study | See Table 1 |
| **Software and algorithms** |  |  |
| GraphPad Prism v9.4.0 | GraphPad Software Inc. | https://www.graphpad.com/ |
| FlowJo v10.8.1 | TreeStar, Ashland, OR, USA | https://www.flowjo.com/ |
| ggplot 2 R package v3.4.0 | Wickham et al. <sup>[82]</sup> | https://ggplot2.tidyverse.org/ |
| Seurat v4.0.4 | Hao and Hao et al. <sup>[83]</sup> | https://satijalab.org/seurat/ |
| Cell Ranger v6.0.1 | Coomes et al. <sup>[84]</sup> | https://www.10xgenomics.com/support/software/cell-ranger/latest |

### Viruses and Tissue Culture

Mouse 3T12 fibroblasts (ATCC CCL-164), mouse J774A.1 macrophages (“J774”; ATCC TIB-67), and mouse RAW264.7 macrophages (ATCC, TIB-71) were cultured in complete DMEM (cDMEM): Dulbecco’s modified Eagle medium (DMEM; Life Technologies) supplemented with 5% and 10% fetal bovine serum (FBS; Atlanta Biologicals) respectively, 2 mM L-glutamine, 10 U/mL penicillin, and 10 μg/mL streptomycin sulfate. Cells were cultured at 37°C with 5% CO_2_. Wild-type (WT) MHV68, MHV68.LANA::βlac, MHV68.H.GFP [53], MHV68.ORF59.GFP [54], MHV68.vCycKO [59], MHV68.CA-RTA [81] and MHV68.vCyc::blac viruses were grown and prepared as described previously [43, 44, 53], with virus titers determined from at least three independent plaque assays. MHV68.vCyc::blac was created through BAC recombineering, introducing a C-terminal fusion with the blac gene downstream of the endogenous vCyc (ORF72) gene (kindly provided by Dr. Darby Oldenburg, unpublished data). All recombinant viruses used in these studies had the BAC origin of replication removed prior to viral stock generation. All recombinant viruses used in these studies are capable of robust lytic replication in fibroblasts, replicating to high-titers of infectious virus with kinetics comparable to WT MHV68 (data not shown) [43, 44, 53, 54, 59, 81].

### Infection

All in vitro infections were based on live cell counts with the TC20 Automated Cell Counter (Bio-Rad) with Trypan Blue dye (Bio-Rad, Cat. No. 145-0021). Viral stock was diluted with cDMEM for viral inoculum with a multiplicity of infection (MOI) of 1 or 10 PFU/cell. Cells were incubated at 37°C with 5% CO_2_ for 1 hour, with rocking every 15 min, before viral inoculum was removed and replaced with 1mL of cDMEM. For qPCR analysis of viral DNA replication and for beta-lactamase staining and flow cytometric analysis of infected cells, viral inoculum was left on cells after 1 hr infection time. For IL-4 treatments, media containing recombinant mouse IL-4 (PeproTech, Cat. No. 214-14) was added to cells 16 hours before infection. Following infection, media containing 0, 10 or 100ng/mL of fresh IL-4 containing media were added respectively.

Samples were harvested at the indicated hours post-infection (hpi) through either cell scraping (for DNA isolation) or through incubation in TRIzol (for RNA isolation).

### Plaque Assay

Plaque assay quantification of viral titer was performed using 3T12 cells. Cells were plated in 12-well plates at 8.5×10^4^ cells per well one day prior to infection. Viral samples were diluted 10-fold in 5% cDMEM. An internal standard was included for each infection to ensure reproducible sensitivity for each plaque assay. Cells were incubated with virus for 1 hour at 37°C with 5% CO_2_. Plates were rocked every 15 min. Cells were then covered with an overlay composed of a 1:1 mix of 10% cDMEM and carboxymethyl cellulose (CMC; Sigma, Cat. No. C-4888) supplemented with Gibco™ Amphotericin B (Thermo Fisher Scientific, Cat. No. 15290018). Cells were incubated for 8 days before staining with 0.5% methylene blue and plaques were counted.

### Western Blot

Cell extracts were lysed using RIPA lysis buffer (50mM TrisCl pH 8.0, 1mM EDTA, 1% Triton X-100, 0.5% Sodium Deoxycholate, 0.1% SDS, 140mM NaCl) and 1X proteinase and phosphatase inhibitor cocktail (Thermo Fisher, Cat. No. PI78440). Protein concentration was measured by Bio-Rad DC Asay (Bio-Rad Cat. No. 5000111) and normalized to 30μg. Samples were mixed 1:1 with 1X Laemmli buffer (Bio-Rad, Cat. No. 1610747) and briefly heated for 5min at 95° C prior to loading samples on a 10% SDS-polyacrylamide gel (Bio-Rad, Cat. No. 4561033). Samples were transferred to a PVDF membrane using a semi-dry transfer, followed by immunoblotting using a mouse monoclonal beta-actin (Thermo Fisher, Cat. No. MA5-15739) and a chicken anti-ORF75C antibody (a gift from Dr. Laurie Krug, NCI/CCR, Bethesda, Maryland, USA) [80]. Blots were visualized using Bio-Rad Clarity ECL Substrate (Bio-Rad, Cat. No. 1705061).

### RT-qPCR

RNA was isolated from infected cells harvested at indicated times by 10-minute incubation in TRIzol® reagent (Thermo Fisher Scientific, Cat. No. 15596026), followed by TURBO™ DNase (Invitrogen, Cat. No. AM2238) treatment according to manufacturer’s protocols. RNA amplification and removal of DNA was confirmed by PCR amplification of the control host gene, 18S, in the presence or absence of reverse transcriptase. RNA presence and absence of DNA was confirmed with RT-PCR and PCR respectively. RT-PCR was performed using the OneStep RT-PCR kit (Qiagen, Cat. No. 210212) with the following conditions: (i) 50°C for 30 min, (ii) 95°C for 15 min, (iii) 40 cycles of 94°C for 30 sec, 52°C for 30 sec, and 72°C for 30 sec, (iv) 72°C for 10 min, and (v) hold at 4°C. PCR was performed using *Taq* DNA polymerase (Qiagen, Cat No. 201205) with the following conditions: (i) 95°C for 5 min, (ii) 40 cycles of 94°C for 30 sec, 52°C for 30 sec, and 72°C for 30 sec, (iii) 72°C for 10 min, and (iv) hold at 4°C. RNA samples that showed no product following PCR amplification were deemed DNA-free, and then converted to cDNA using random primers (250ng/uL) (Invitrogen, Cat. No. 48190011) and SuperScript II reverse transcriptase (Invitrogen, Cat. No. 18064014) following the manufacturer’s protocol. 100 nanograms of cDNA was used for qPCR analysis of the identified genes (Quantitect Primer Assay, Qiagen) using the iQ SYBR green supermix (BioRad Cat. No.1708880) with the following conditions: (i) 95°C for 3 min, (ii) 40 cycles of 95°C for 15 sec, 60°C for 1 min, and (iii) 95°C for 15 sec, 60°C for 1 min, and 95°C for 15 sec. Amplification of viral genes was normalized to murine 18S expression to calculate the relative difference of target gene expression using the Pfaffl method, as previously described [6, 85] : Target primer efficiency^TargetL1Ct/18S primer efficiency^18SL1Ct. A single product for each target was confirmed by melt curve analysis. PCR primers are listed in **Table 1**.

**Table 1:**
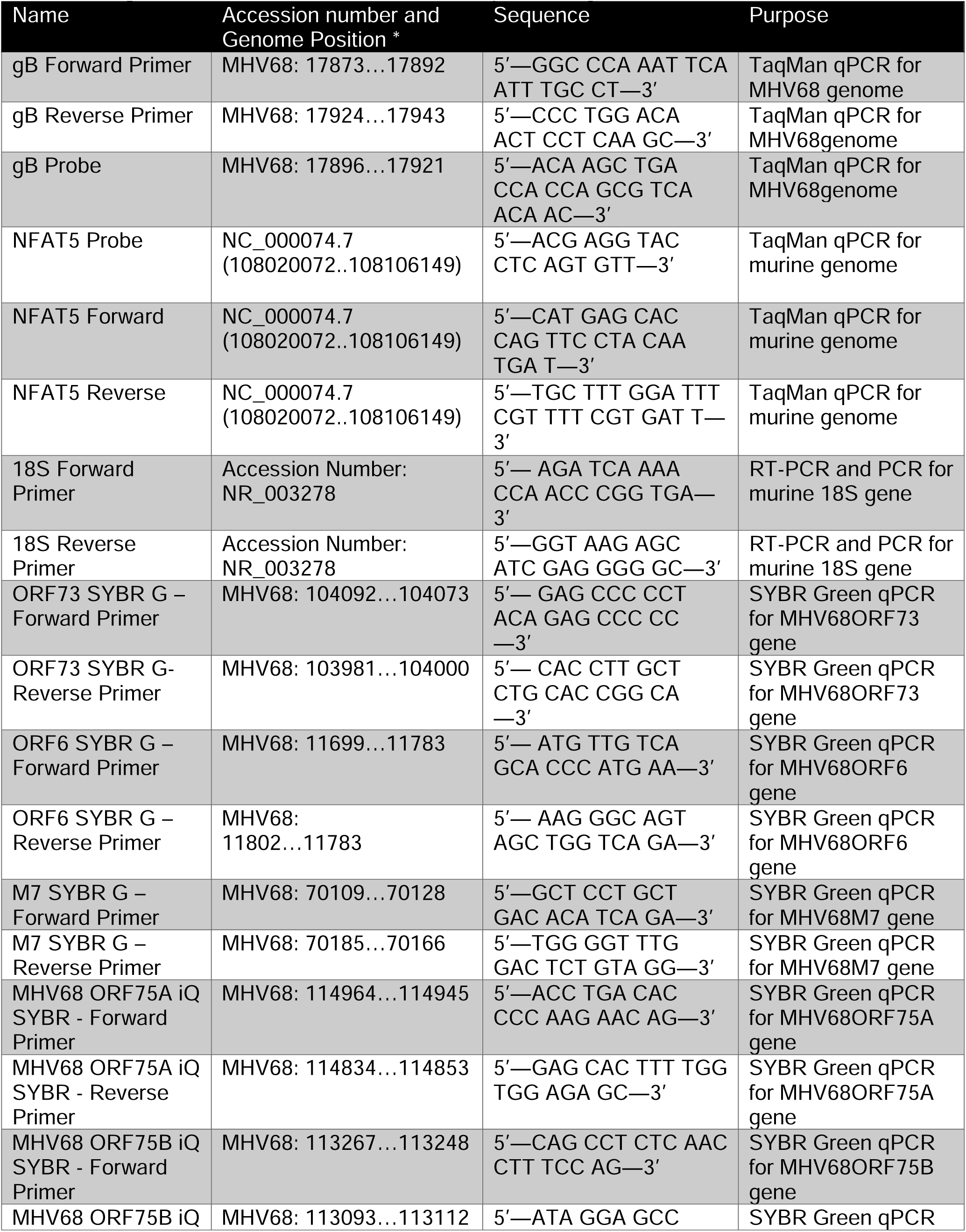
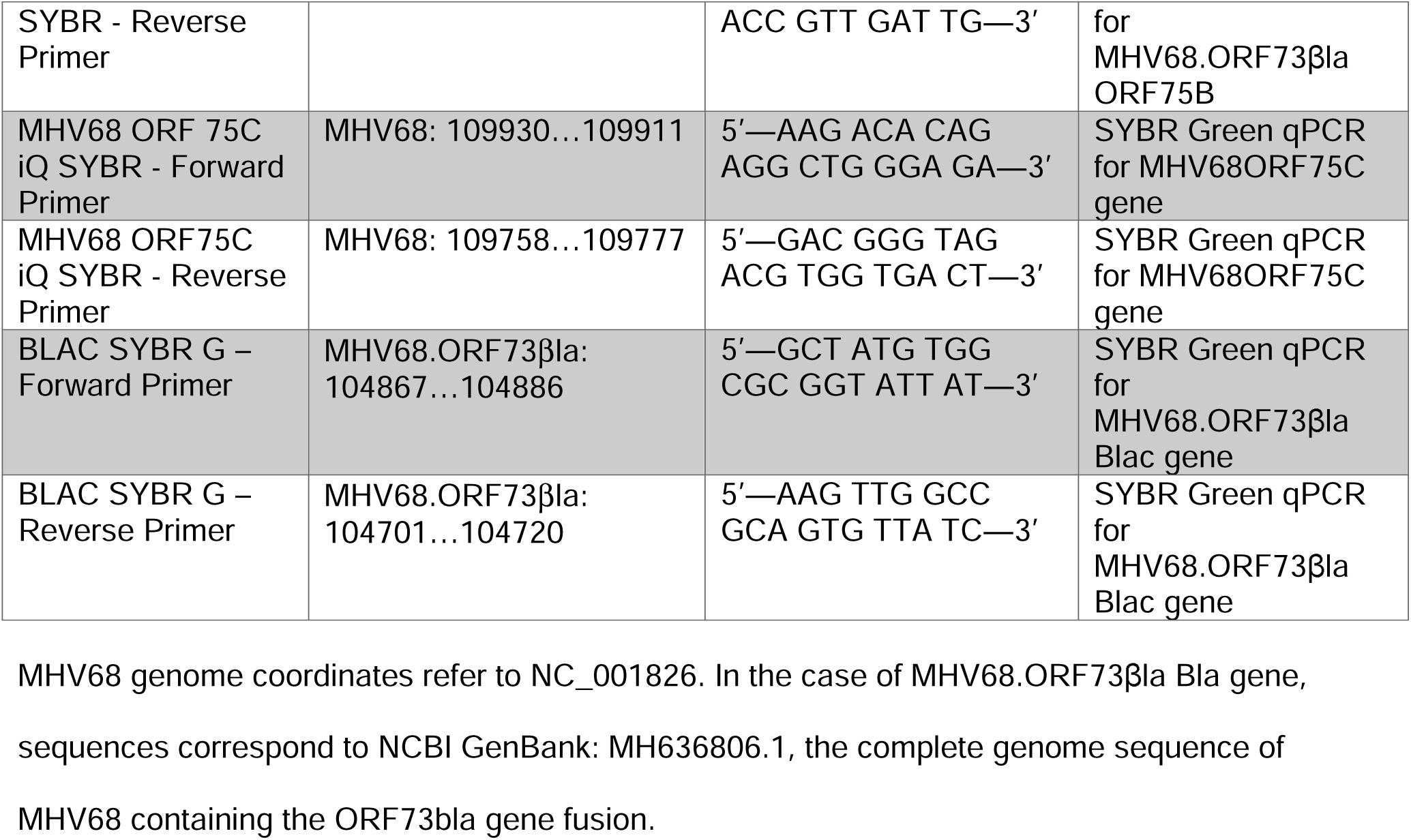
Oligonucleotides for RT-PCR, PCR, and qPCR analysis.

### Quantitative PCR for quantification of viral DNA replication

Infected cells were harvested by cell scraping at time (hpi) indicated. Harvested cells underwent three freeze/thaw cycles and DNA was isolated using the DNeasy Blood and Tissue Kit (Qiagen, Cat. No. 69504), with an overnight proteinase K incubation and heat inactivation at 56°C. DNA was normalized to a concentration of 20 ng/uL in molecular grade water. qPCR analysis was done on 100ng of DNA using a LightCycler 480 Probe Master-Mix kit (Roche, Cat. No. 04707494001) and a primer and probe set specific to MHV68 glycoprotein B (gB) to quantify the number of viral genome copies. Viral genome copies/cell concentration was determined by dividing gB quantity over NFAT5 quantity. Primers and probes listed in Table 1. A gB or nuclear factor of activated T cells 5 (NFAT5) [84, 86] standard curve was generated using a gB or NFAT5 plasmid dilution series ranging from 10^2^ to 10^10^ copies diluted in background DNA, with a limit of detection (LOD) of 100 copies.

### Single-cell RNA Sequencing

To characterize MHV68 transcription during primary, acute peritoneal macrophage infection *in vivo*, female C57BL/6 mice were infected with 1x10^6^ PFU by intraperitoneal injection. Peritoneal cells were harvested and pooled from four infected animals at 16 hours post-infection, stained with the live/dead discrimination dye Zombie Near-IR (Zombie NIR^TM^, BioLegend, Cat. No. 423105), F4/80-AlexaFluor 674 (clone BM8, 1:200 dilution) in the presence of Fc receptor blockade (clone 2.4G2), and CCF2-AM (Thermo Fisher, Cat. No. K1023). Cells were sorted on a MoFlo Astrios, purifying live, single, F4/80+ cells that either had CCF2 cleavage (indicating virus infection, defined by expression of the LANA::blac fusion protein) or lacking CCF2 cleavage (indicating cells that were not infected or failed to express the LANA::blac fusion protein). Single cell RNA-seq was prepared using the Chromium 3’ V3 library kit (10X Genomics). Cell processing and sequencing was performed by the University of Colorado Cancer Center Genomics Shared Resource (RRID: SCR_021984). Resulting data were processed as follows: Cell Ranger (v6.0.1) [87] was used to process the fastq files to cell and gene count tables using unique molecule identifiers (UMIs) with the include-introns parameter. Because of the difficulties counting viral reads, this was performed in a two-pass manner. In the first pass, reads were aligned to a chimeric genome of mouse mm10 (GENCODE M23 gene annotations) and MH636806.1 (no gene annotations, only positive and negative strand alignment). The viral counts were stored in metadata and removed from the counts matrix. Reads mapping to MH636806.1 in step 1 were aligned to the same chimeric genome but in this case, the viral transcriptome as well as specific intergenic regions were annotated. The intergenic regions were determined as the strand-specific sequences between genes with 1 base pair of padding on both ends. To minimize reads overlapping with multiple viral genes, only the first 70bp of each read were aligned and counted. The resulting counts matrix of viral alignment was appended to the host gene counts from step 1.

The Seurat (v4.0.4) [83] pipeline was used for downstream quality control and analysis.

Cellranger-filtered data was read into Seurat. Host genes were removed if identified in fewer than 10 cells while viral gene and intergenic regions were removed if found in no cells. Cells were removed if they expressed < 50 genes or viral regions, < 5000 UMIs, > 5 % total UMIs from mitochondria, or > 70000 UMIs. The filtered data was normalized by dividing gene counts by total counts per cell and multiplied by 10,000 followed by natural-log transformation. The top 2,000 most variable genes were scaled with total UMI and percentage mitochondria regressed out. These were used to calculate principal components (PCs) with the top 20 used to perform Uniform Manifold Approximation and Projection (UMAP) and determining the k-nearest neighbors and clustering.

Clusters were identified by over-representation analysis and examining the top enriched markers. Canonical cell type markers were plotting in UMAP space and using dot plots. Dot plots, bar plots, scatter plots, and violin plots were generated using Seurat and ggplot2 R packages (v3.4.0) [82].

### Flow Cytometry

3T12 and J774 cells were infected with MHV68 LANAβlac or MHV68.H.GFP at an MOI of 1 PFU/cell and harvested at the indicated time points followed by staining for flow cytometric analysis. Samples treated with the recombinant murine IL-4 were given 10% cDMEM with 10ng/mL of IL-4 eighteen hours prior to infection and replaced with fresh media containing IL-4 immediately following infection. Cell suspensions were stained with LIVE/DEAD fixable Near-IR Dead Cell stain kit (Invitrogen, Cat. No. L10119) at a 1:1000 dilution in BSS wash (111mM Dextrose, 2mM KH_2_PO_4,_ 10mM Na_2_HPO_4,_ 25.8 CaCl_2_•2H_2_O, 2.7mM KCl, 137mM NaCl, 19.7mM MgCl•6H_2_O, 16.6mM MgSO_4_). Cells were either incubated with CCF2 or antibody stained for protein detection. Cells for CCF2 detection were stained with CCF2-AM (Thermo Fisher, Cat. No. K1023) at a 1:10 dilution following the manufacturer’s protocol. To detect proteins, cells were stained with Live/Dead Fixable Blue (Thermo Fisher, Cat. No. L23105) at a 1:1000 dilution in BSS wash, Fc receptor block (Fc Shield, Tonbo Biosciences, Cat. No. 70-0161), a rabbit antibody against the MHV68 ORF4 protein (vRCA, 1:400 dilution) (gift of the Virgin Lab, Washington University St. Louis [6, 79]) labeled with a Zenon R-phycoerythrin anti-rabbit IgG reagent (Invitrogen, Cat. No. Z25355), per manufacturer’s recommendations, and a AF647-conjugated anti-mouse phospho-histone H2AX antibody (1:800 dilution) (Sigma Aldrich, Cat. No. 07-164-AF647, clone JBW30). Samples were fixed in 1% paraformaldehyde prior to analysis using the Bio-Rad ZE5/YETI Cell Analyzer or an Agilent Novocyte Penteon flow cytometer. To detect actin mRNA, cells were stained with buffers and probes using the PrimeFlow RNA assay kit (ThermoFisher), as previously published [6]. All flow cytometry experiments included unstained, single-stain and full minus one controls to define background fluorescence, fluorescent signal spread and compensation.

### Light microscopy

At the indicated time points, 6-well plates of 3T12 or J774 cells were imaged on a Nikon Eclipse Ti2 inverted light microscope equipped with Iris 15 camera (Photometris) and use of NIS elements software. Plates were mounted on a stage heated to 37°C. A 10x phase objective was used to capture brightfield images. Immediately following image acquisition, samples were harvested and processed for flow cytometric analysis.

### Statistical Analysis and Software

Data analysis, graphing and statistical analyses were performed using GraphPad Prism (version 9.4.0; GraphPad Software, San Diego, California USA, www.graphpad.com). Flow cytometry data were analyzed using FlowJo (version 10.8.1. Ashland, OR: Becton, Dickinson and Company; 2022). scRNA-seq data processing and analysis were done as described above. Statistical significance was tested by unpaired, nonparametric, Mann-Whitney t test or 1-way ANOVA for comparing replicate values from experiments. Raw and processed single cell RNA-Seq data is available through NCBI GEO (GSE233202). Statistical analysis tests and statistical significance are defined in figure legends.

### Cell Sorting

J774 fibroblasts were infected at an MOI=1 with for one hour, after which inoculum was removed. Cells were harvested at 48 hours post-infection (hpi) and stained with β-lactamase substrate as previously reported [44] using the CCF2-AM substrate (Invitrogen, Life Technologies). Cells were subjected to fluorescence activated cell sorting (FACS), with cells gated on singlet cells that were positive for the cleaved CCF2-AM substrate (subsequently referred to LANAβlac+ sorted cells). Cleavage of CCF2-AM was defined as positive fluorescence in the 405-448/59nm emission channel obtained in WT MHV68.LANAβlac, compared to parental WT MHV68 infected cultures with no beta-lactamase fusion gene, used to define background fluorescence. β-lactamase positive cells were sorted using a Astrios EQ (Beckham Coulter). Cells were either plated onto 6-well plates at 5 x 10^5^ cells/mL, collected for quantitative PCR or plaque assay measurement. The plated CCF2+ J774 cells were cultured at 37°C with 5% CO_2_, grown in complete DMEM (cDMEM): Dulbecco’s modified Eagle medium (DMEM; Life Technologies) supplemented with 10% fetal bovine serum and passaged upon confluency (typically every 3-4 days). Upon passaging, cells and supernatant were collected followed by resuspension in 1mL of 10% cDMEM. Cell counts were measured by BIORAD TC20 automated cell counter. At 7 days post sort (dps), one 6-well of cells was collected and resuspended in 1mL of 10% cDMEM and flow cytometry for CCF2 detection was performed with 700uL of the resuspended cells. At 14 dps, one 6-well of cells was collected and resuspended in 1mL of 10% cDMEM and flow cytometry for CCF2 detection and viral protein vRCA and γH2AX was performed with 700uL of the resuspended cells (See Flow Cytometry Methods Section). At both 7 and 14 dps 300uL of the 1mL collected cells were used for qPCR and plaque assay analysis.

### Ethics Statement

All animal studies were performed in accordance with the recommendations in the Guide for the Care and Use of Laboratory Animals of the National Institutes of Health. Studies were conducted in accordance with the University of Colorado Denver/Anschutz Institutional Animal Use and Care Committee under the Animal Welfare Assurance of Compliance policy (assurance no. D16-00171). All procedures were performed under isoflurane anesthesia, and all efforts were made to minimize suffering.

## Acknowledgments

We thank Dr. Brian Russo’s lab, especially Jenna Vickery for the use and training of their Nikon Eclipse Ti2 inverted light microscope, and Christine Childs, Kristina Terrell and Dmitry Baturin of the University of Colorado Cancer Center Flow Cytometry Shared Resource for training and assistance with flow cytometry. We thank Dr. Laurie Krug and Dr. Craig Forrest for generously sharing the ORF59.GFP recombinant virus. R.E.K. was supported by the Molecular Biology T32 pre-doctoral training grant (NIH 5T32GM136444). C.G. was supported by the University of Colorado School of Medicine RNA Bioscience Initiative Summer Internship Program. This study was supported by National Institutes of Health grant R01 AI157201 awarded to LvD and ETC, with additional support by the National Institutes of Health Cancer Center Support Grant (P30CA046934), including assistance from the Bioinformatics and Biostatistics Shared Resource (RRID: SCR_021983), Flow Cytometry Shared Resource (RRID: SCR_022035), and Genomics Shared Resource (RRID: SCR_021984). Generative AI was not used in preparation of this manuscript.

## Contributor Roles

Conceptualization: G.V., E.T.C., L.F.v.D. Methodology: D.G.O.

Validation: G.V., K.S.N., A.T., R.E.K., E.A.A., S.C., C.G., M.D., B.D.G.

Formal analysis: A.G., E.T.C.

Investigation: G.V., K.S.N., A.T., R.E.K., E.A.A., S.C., C.G., M.D., B.D.G., E.M.M.

Data curation: G.V., A.G., E.T.C.

Writing – original draft: G.V., E.T.C., L.F.v.D.

Writing – review & editing: G.V., E.T.C., L.F.v.D.

Visualization: G.V., A.G., E.T.C., L.F.v.D. Supervision: E.T.C., L.F.v.D.

Project administration: G.V., E.T.C., L.F.v.D. Funding acquisition: E.T.C., L.F.v.D.

---

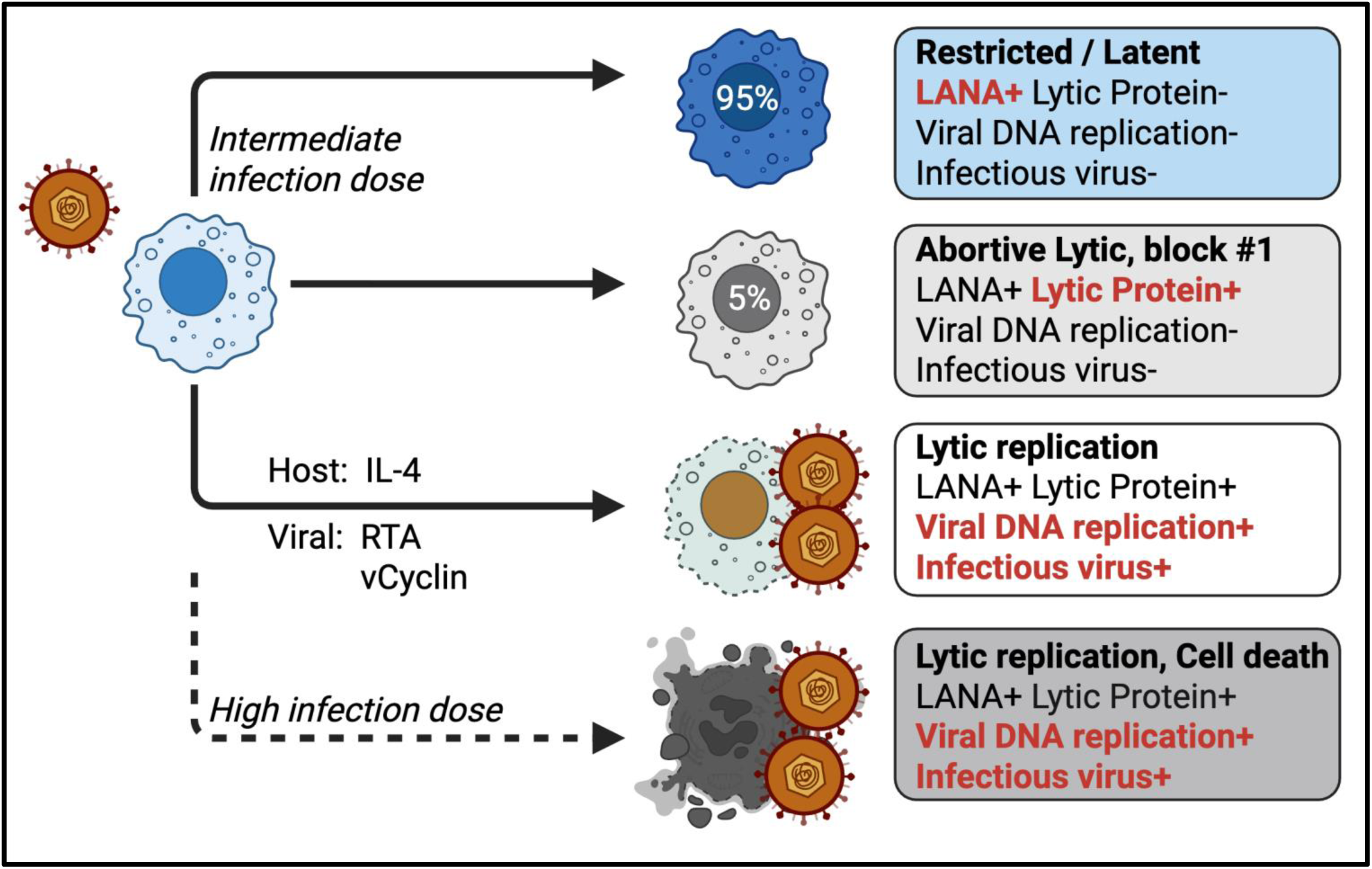

- MHV68 efficiently infects macrophages
- Infection results in three divergent outcomes regulated by discrete inputs
- Macrophages can support lytic replication, promoted by the host cytokine IL -4 and viral genes.
- High infection dose is associated with a significantly increased frequency of cell death.

**Supplemental Figure 1 (Figure S1).**
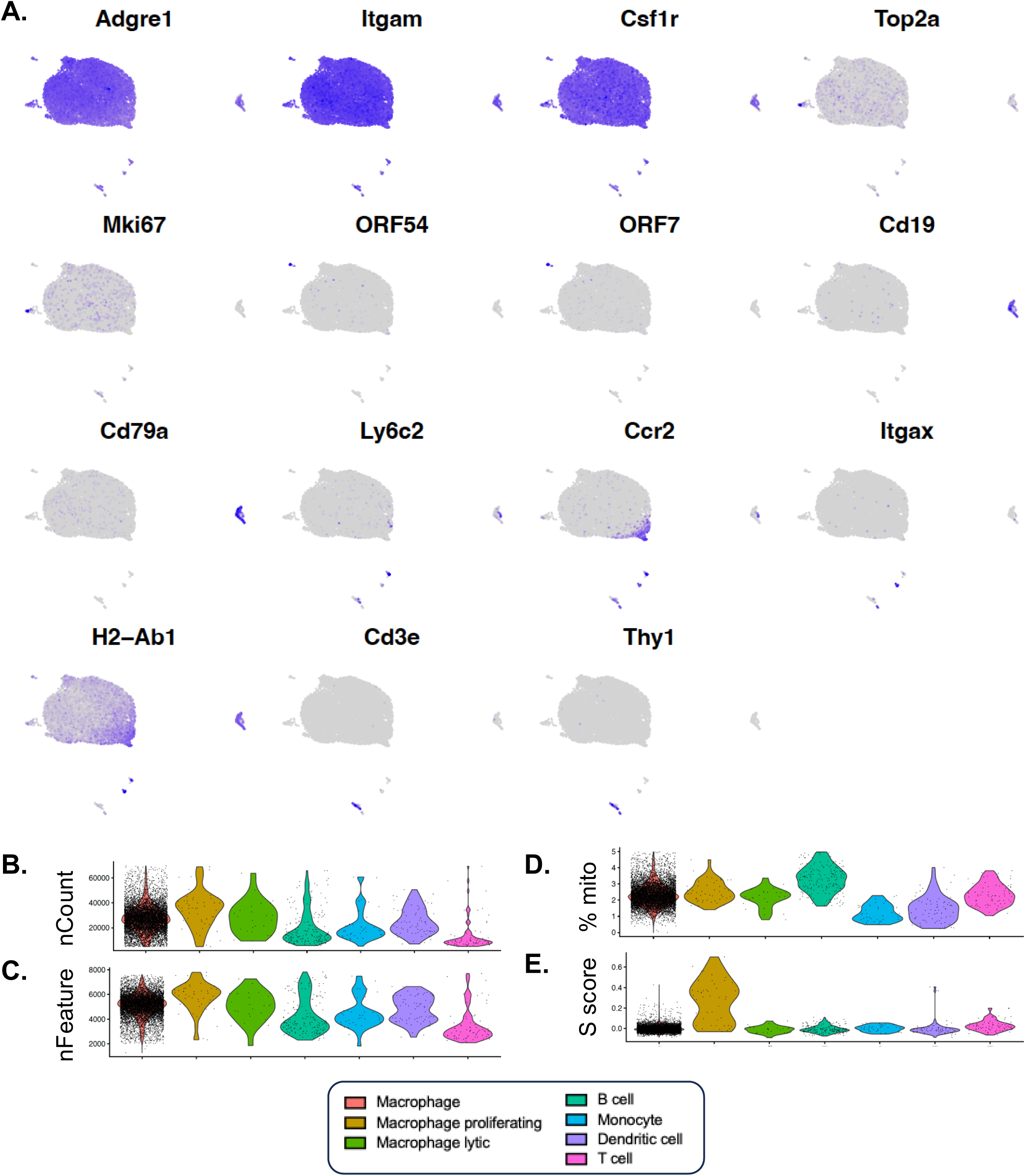
Single cell RNA sequencing analysis of acute MHV68 macrophage infection. Data correspond to **Figure 1**. (A) Visualization of gene expression based on UMAP topography in Fig. 1A. Maximal expression is designated by purple, with absence of detection designated by gray. Each panel depicts a distinct lineage-defining gene used to identify cell types in Fig. 1, with signature genes for each cell type as follows. Macrophage: Adgre1 (encodes the F4/80 protein), Itgam (encodes the CD11b protein). Macrophage proliferative: Top2a, Mki67. Macrophage lytic: ORF54, ORF7. B cell: Cd19, CD79a. Monocyte: Ly6c2, Ccr2. Dendritic cell: Itgax, H2-Ab1. T cell: Cd3d, Thy1. (B-E) scRNA-seq quality control metrics across cell subsets, including (B) UMI Count, (C) Features, (D) % mitochondrial reads, and (E) S score, a proliferation-associated gene signature.

**Figure S2.**
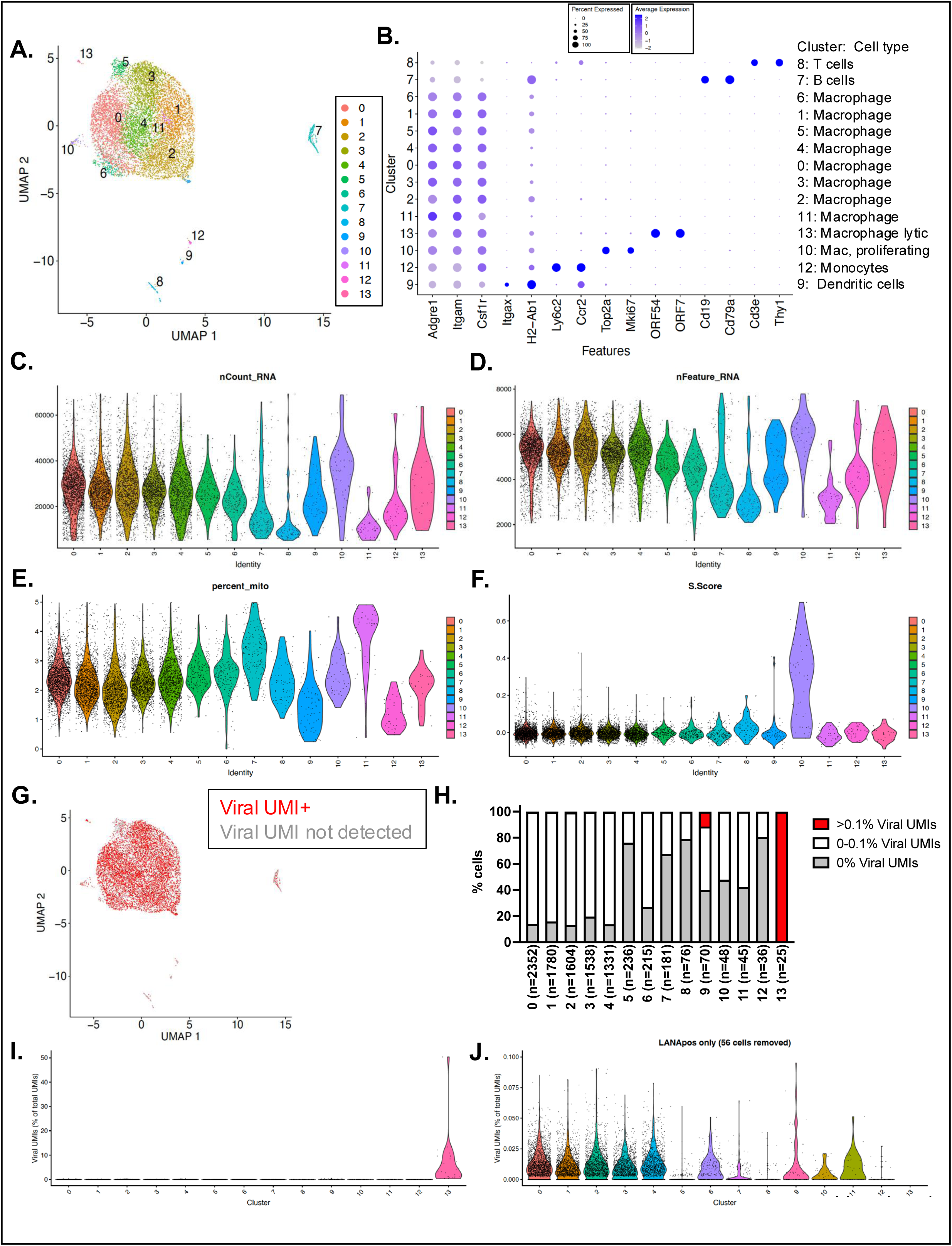
Single cell RNA sequencing analysis of acute MHV68 macrophage infection analyzed using a higher number of cell clusters. Data correspond to **Figure 1**, with scRNA-seq subjected to Seurat-based designation of 14 clusters. **(A)** UMAP visualization of Seurat-defined clusters. **(B)** Visualization of the frequency and magnitude of host gene expression across clusters, with cell type designation indicated on right. **(C-F)** Quality control metrics across the 14 clusters, including (C) UMI Count, (D) Features, (E) % mitochondrial reads, and (F) S score, a proliferation-associated gene signature. **(G)** UMAP visualization of cells with at least 1 detectable viral UMI per cell identified in red. **(H)** Frequency of cell subsets stratified by the frequency of viral UMIs of total UMIs detected per cell. **(I-J)** Percentage of viral UMIs out of total UMIs across clusters (panel I, y axis range from 0-50%), or (J) focused only on the majority of virus positive cells (y axis range, 0- 0.1%; excluding 56 cells with % viral UMI > 0.1). Data are from 9,537 cells, with clusters containing between 25-2,532 cells. Panel G is included to provide a frame of reference and depicts the same data as in Figure 1D.

**Figure S3.**
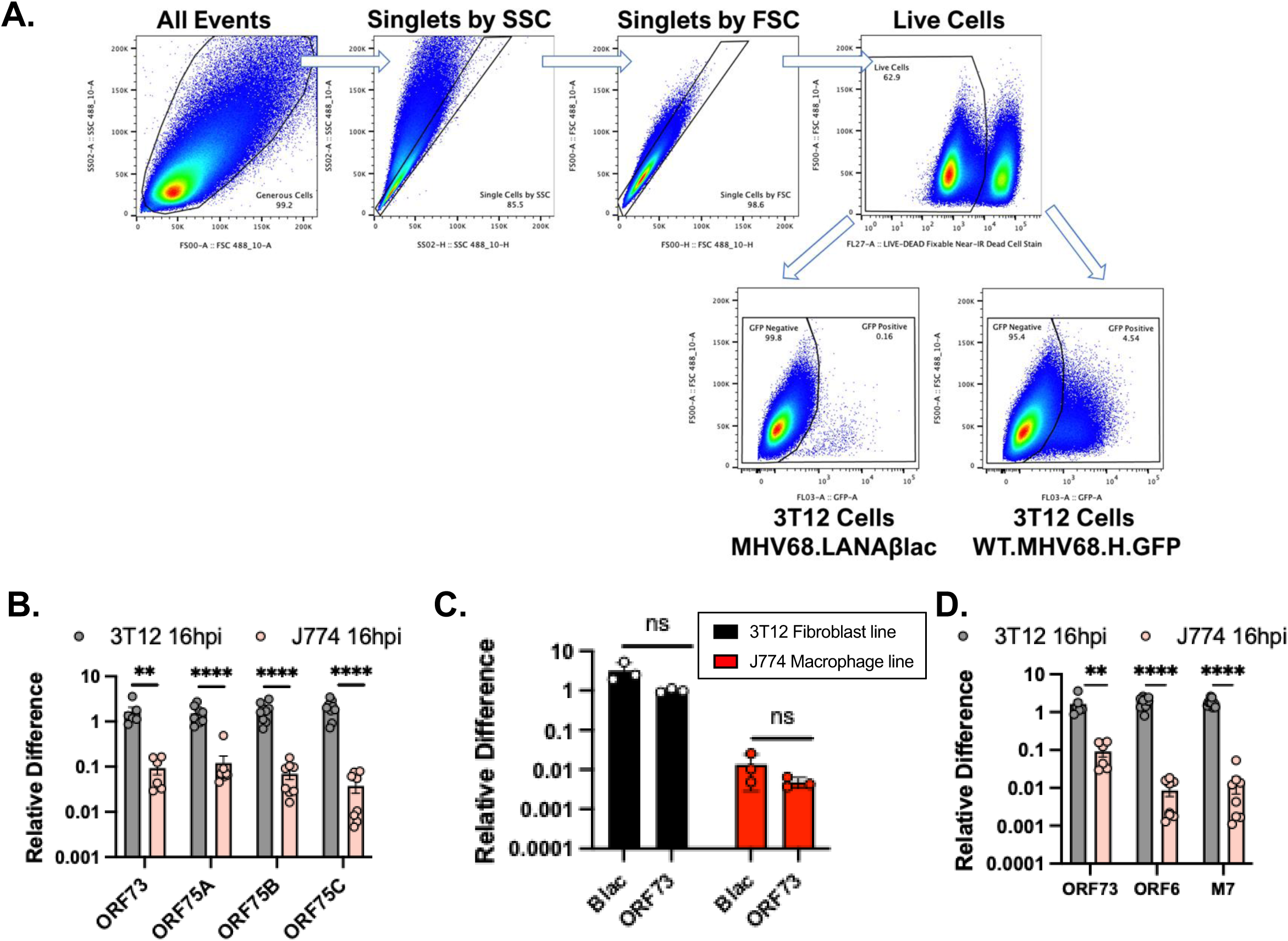
Flow cytometry gating strategy and viral gene expression during MHV68 infection. **(A)** Flow cytometry gating strategy to analyze MHV68-infected cells, in support of **Figure 3E**. 3T12 cells infected with MHV68 LANAblac or WT.MHV68.H.GFP were analyzed by flow cytometry, using sequential focused analysis (i.e. ”gating”) to consider cells that are single cells (defined sequentially as “Singlets by SSC” and “Singlets by FSC”) that are viable (”Live cells”) defined by exclusion of a dye that identifies dead cells. Live cells were then analyzed for GFP fluorescence as measured by the GFP detection channel. Cells infected with MHV68.LANAβlac, without additional of the CCF2 fluorescent substrate, identified background fluorescence, in contrast to samples infected with MHV68.H.GFP which encoded GFP. Controls included single-stained samples with only one stain to define the bounds of a gate for the fluorophore of interest. **(B-D)** Viral gene expression defined by qRT-PCR in 3T12 or J774 cells infected with WT MHV68.LANAblac (MOI=1) at 16 hpi (B,D) or 24 hpi (C) in support of **Figure 3F-G**. Data show mean ± SEM from 3 independent experiments with 3 biological replicates measured in technical triplicates per experiment (depicted by individual symbols) (B,D) or from 1 experiment with 3 biological replicates measured in technical triplicates, with ORF73 data from Fig. 2B (C). Statistical analysis was done using unpaired t test with statistically significant differences as indicated. ** p<0.01, ***p<0.001, ****p<0.0001. ns, no statistical significance detected.

**Figure S4.**
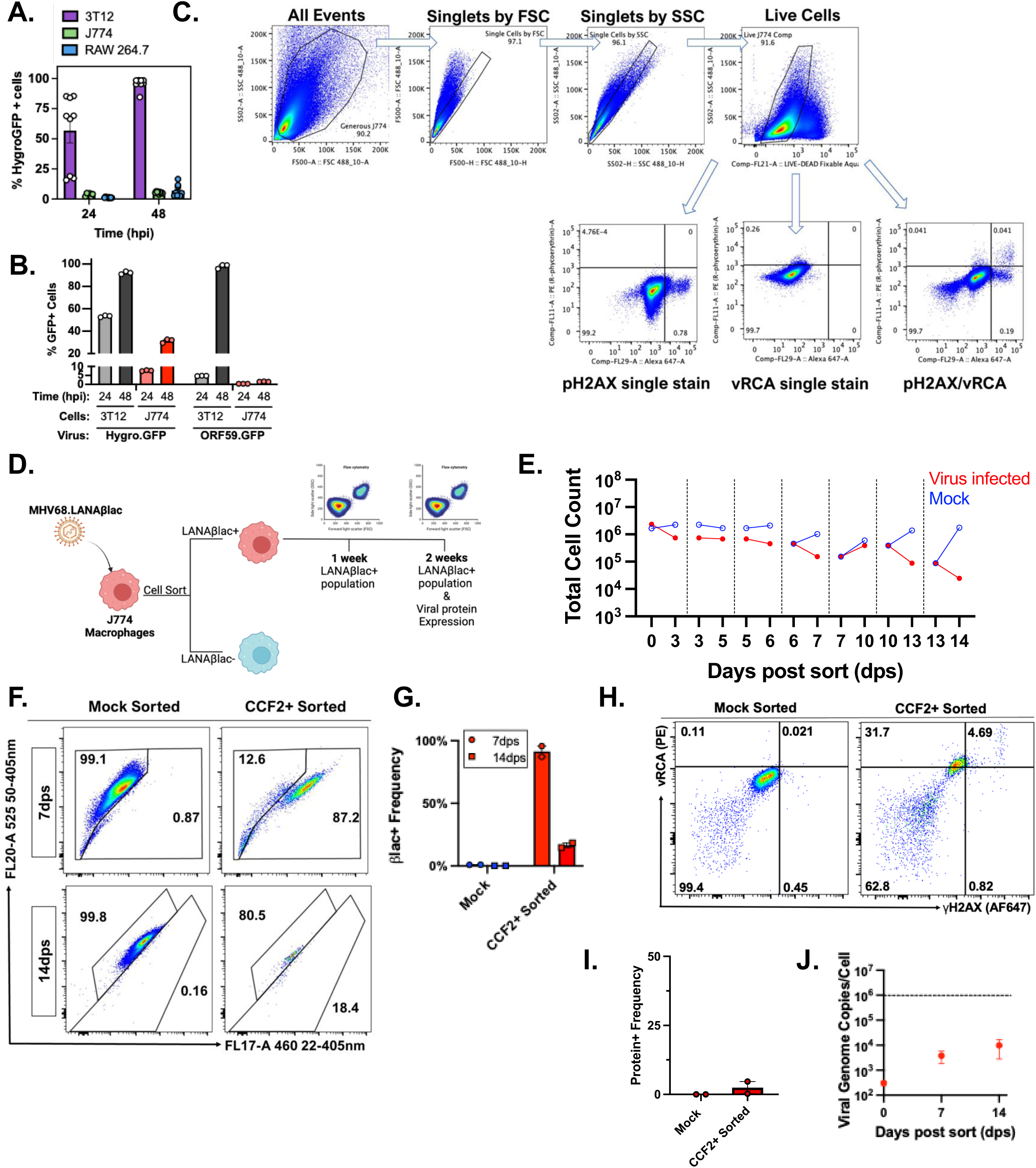
Gating of flow cytometry analysis of MHV68 LANAβlac infected J774 cells for lytic protein markers. **(A)** Analysis of GFP expression in 3T12 fibroblasts, J774 and RAW264.7 macrophage cell lines at 24 or 48 hpi, following infection with MHV68.H.GFP (MOI=1). **(B)** Gating strategy to analyze vRCA and γH2AX expression in MHV68-infected J774 cells, in support of **Figure 4C**. Cells were analyzed by flow cytometry, using sequential focused analysis (i.e. ”gating”) to consider cells that are single cells (defined sequentially as “Singlets by SSC” and “Singlets by FSC”) that are viable (”Live cells”) defined by exclusion of a dye that identifies dead cells. Live cells were then analyzed for antibody binding for lytic protein markers. Gate boundaries were defined using a combination of “single stain” samples, containing only the fluorophore of interest, and “full minus one” controls, lacking the antibody in the fluorescent channel of the excluded fluorophore (not shown). **(C-I)** Analysis of MHV68-infected, FACS-purified J774 cells over time. **(C)** Experimental overview with J774 macrophages infected with MHV68.LANAβlac (MOI=1), followed by sorting for LANAblac+ cells defined based on CCF2 substrate cleavage. Mock and LANAblac+ sorted cells were cultured and analyzed over 14 days. **(D)** Total cell counts over the course of the study, with each vertical line indicate a subsequent cell passage. **(E)** Flow cytometric analysis of LANAβlac expression defined by cleavage of the CCF2 fluorescent beta-lactamase substrate, analyzing single, viable cells, with values in the right gate defining the frequency of cells with CCF2 substrate cleavage. **(F)** Quantification of LANAβlac+ cell frequency defined by flow cytometry, as in panel E. **(G)** Flow cytometric analysis of MHV68 lytic cycle-associated proteins (vRCA, γH2AX) at 14 dps, comparing mock (left) or CCF2-sorted, LANAblac+ cells (right). Data depict events that were single, viable cells, with values defining the frequency of cells with vRCA and/or phosphorylated host γH2AX, indicative of lytic MHV68 infection, as in panel B. **(H)** Frequency of vRCA+ γH2AX+ cells defined by flow cytometry, as in panel G. **(I)** Quantitation of viral DNA by qPCR. Dotted line indicates mean 3T12 viral genome copies/cell at 72 hpi. Data show mean ± SEM from 3 independent experiments with 3 biological replicates per experiment (depicted by individual symbols) (A) or from 2 independent experiments.

## Notes

### Competing Interest Statement

The authors have declared no competing interest.

### Summary of Updates

Updated data on Figure 5 (H, I) and associated methods. Clarifications in text.

